# A Cellular Basis for Mapping Behavioural Structure

**DOI:** 10.1101/2023.11.04.565609

**Authors:** Mohamady El-Gaby, Adam Loyd Harris, James C. R. Whittington, William Dorrell, Arya Bhomick, Mark E. Walton, Thomas Akam, Tim E. J. Behrens

## Abstract

To flexibly adapt to new situations, our brains must understand the regularities in the world, but also in our own patterns of behaviour. A wealth of findings is beginning to reveal the algorithms we use to map the outside world^1–6^. In contrast, the biological algorithms that map the complex structured behaviours we compose to reach our goals remain enigmatic. Here we reveal a neuronal implementation of an algorithm for mapping abstract behavioural structure and transferring it to new scenarios. We trained mice on many tasks which shared a common structure organising a sequence of goals, but differed in the specific goal locations. Animals discovered the underlying task structure, enabling zero-shot inferences on the first trial of new tasks. The activity of most neurons in the medial Frontal cortex tiled progress-to-goal, akin to how place cells map physical space. These “goal-progress cells” generalised, stretching and compressing their tiling to accommodate different goal distances. In contrast, progress along the overall sequence of goals was not encoded explicitly. Instead a subset of goal-progress cells was further tuned such that individual neurons fired with a fixed task-lag from a particular behavioural step. Together these cells implemented an algorithm that instantaneously encoded the entire sequence of future behavioural steps, and whose dynamics automatically retrieved the appropriate action at each step. These dynamics mirrored the abstract task structure both on-task and during offline sleep. Our findings suggest that goal-progress cells in the medial frontal cortex may be elemental building blocks of schemata that can be sculpted to represent complex behavioural structures.

## Introduction

Our behaviours are highly structured. From cooking a meal to solving a maths problem, we compose elaborate sequences of actions to achieve our goals. When elements of this structure are common across tasks, we can build a schema; a generalised representation of task states that allows us to instantly compose new behavioural sequences, and infer steps never taken before^7,8^. These behavioural structures can be complex and hierarchical, where sequences of actions leading to individual goals are nested within a higher order structure relating the goals to each other^9,10^. To successfully execute a hierarchical behavioural sequence, one must simultaneously track their position at all levels of the task hierarchy.

Lesion, imaging and neurophysiological studies implicate the frontal areas of the neocortex in mapping task structure. This involves roles in forming a schema of task structure^11–17^, generating complex behavioural sequences^18,19^, encoding goals^20,21^ and simultaneously tracking a working memory of multiple task variables^22–24^. These findings are consistent with a role of the frontal cortex in generating a map of abstract task structure: a representation that reflects the conserved relationships between task states and generalises across similarly structured tasks with different sensorimotor specifics. A key remaining challenge is to derive biological algorithms that mechanistically explain how frontal activity generates such abstract maps.

What mechanisms mediate the mapping of abstract task structure? Such mapping should comprise neuronal dynamics that evolve as a function of progress in a task, rather than related variables such as elapsed time or the number of actions taken. This would naturally enable representations to generalise across goal-directed behavioural sequences that differ in length and duration. Frontal neurons have been found to track progress relative to individual goals^25–27^, regardless of the location of the goal or the distance covered to reach it^27^. However, behavioural tasks are complex and often composed of multiple, hierarchically organised goals. It remains unclear how neurons track progress along such complex tasks. One way this can be achieved is via neurons that invariantly track progress in a sequence of goals regardless of their sensorimotor specifics, in direct analogy to the invariant tracking of progress towards individual goals. Indeed, some findings suggest that frontal neurons are tuned to general task states rather than specific stimuli or actions^11,28–31^. This has led to the view that, in each new task, neurons encoding abstract task states are flexibly bound via synaptic plasticity to those representing the detailed behavioural sequences to be executed^31–34^. Alternatively, a separate line of work on recurrent neural networks suggests that such binding is not necessary and schematic inferences can be made in new scenarios with no new plasticity. Here, details of new task examples are stored as patterns of neural activity using network dynamics sculpted, through the learning of previous examples, by the abstract structure of the task^35^. While the mechanistic details of how such dynamics support task-schema formation remain unclear, this strategy relies on representations that track memories of previous actions and rewards^22–24,35^. Whether and how such representational logic relates to generating a schema that tracks an animal’s progress in task-space remains an open question.

Here we sought to elucidate a neuronal algorithm for mapping abstract task structure. We trained mice on a series of tasks, each of which required visiting 4 goal locations in a repeating, loop-like sequence. The sequential loop structure relating the goals remained the same across tasks, while the goal locations, and hence the behavioural sequences needed to navigate between them, changed. Mice used this abstract structure to perform zero-shot inferences on the first trial of new tasks. Using multi-unit silicon probe recordings, we found that neurons in the medial Frontal Cortex (mFC) tracked progress to the next goal, regardless of the behavioural sequences used to reach it. Crucially, these neurons were further sculpted by the task structure. Neurons were arranged into modules, where activity along each module tracked progress in the higher order sequence of goals. Individual neurons on a given module had mnemonic fields at a fixed lag from a particular behavioural step. Thus, a module behaved like a memory buffer, creating dynamics that track progress in task-space from a particular behavioural step. Each of these memory buffers was shaped by the abstract task structure, reflecting the 4-reward loop, and hence allowed predicting the animals’ future actions a long time before they were made. These findings point to an algorithm that uses <u>S</u>tructured <u>M</u>emory <u>B</u>uffers (SMBs) to encode new behavioural sequences into the dynamics of neural activity without needing associative binding. More broadly, we propose that a schema mapping any complex behavioural structure can be generated by sculpting task-naive progress-to-goal tuning to represent task-structured memories of individual behavioural steps.

## Results

### The ABCD paradigm: an abstract task structure guides rapid sequence learning and inference

We developed a task wherein goal-directed behavioural sequences are hierarchically organised by an abstract structure (the “ABCD” paradigm). Mice (N=11) learned to navigate to identical water rewards arranged in a sequence of 4 locations (***a,b,c*** and ***d***) on a 3×3 grid maze (Figure 1a,b). The reward at each location only became available after the previous goal was visited, so the goals had to be visited in sequence to obtain rewards. Once reward ***d*** was obtained, reward ***a*** became available again, allowing the animal to complete another loop. Each rewarded location (***a,b,c*** or ***d***) defined the beginning of a task “state” (A,B C or D respectively; Figure 1a) and a single ABCD loop defined a trial of a task. A brief tone was played upon reward delivery in state A, marking the beginning of a loop on every trial. Animals encountered multiple tasks where the reward locations changed but the general ABCD loop structure remained the same (Figure 1a). Crucially, task structure was made orthogonal to the structure of physical space: the physical distance between two rewards on the maze was not correlated with their task-space distance (Figure 1a,b, Extended data figure 1a). The task therefore encouraged animals to separately learn the spatial sequences leading to individual goals, and the higher order task structure that organises these goal-directed sequences (Figure 1b).

**Figure 1 –.**
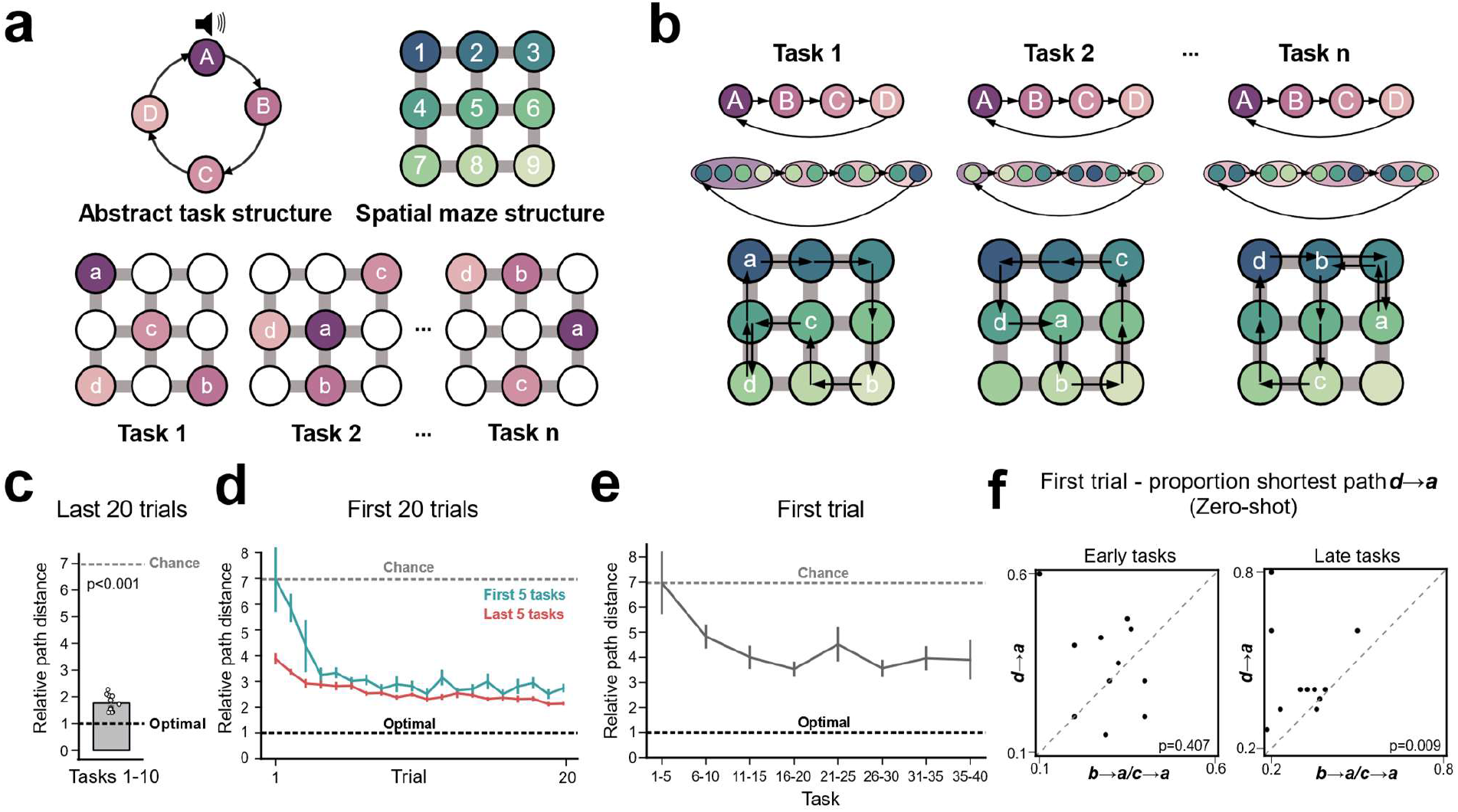
Mice Learn an abstract task structure. a) Task design: animals learned to navigate between 4 sequential goals on a 3×3 spatial grid-maze. Reward locations changed across tasks but the abstract structure, 4 rewards arranged in an ABCD loop, remained the same. b) Learning involved finding optimal sequential paths between each goal. Optimal paths differed in length both within and across tasks but were always organised by an ABCD loop relating all 4 goals. c) When allowed to learn across multiple sessions, animals readily reached near-optimal performance in the last 20 trials, as demonstrated by comparing path length between goals to the shortest possible path (i.e. computing a “relative path distance” measure). T-test against chance (6.96): N=11 mice, statistic=−56.6 P=7.15×10−14 df=10. Chance level was calculated empirically using the mean relative path distance across the first trial of the first 5 tasks. d) Performance improved across the initial 20 trials of each new task. This improvement was markedly more rapid for the last 5 tasks compared to the first 5 tasks. A two-way repeated-measures ANOVA (N=11 mice) showed a main effect of Trial F=11.7 P=6.2×10^−5^, df1=19, df2=190, Task F=26.8 P=4.2×10^−5^, df1=1, df2=10 and a Trial × Task interaction F=3.56 P=0.0201, df1=19, df2=190. e) Performance on the very first trial improved markedly across tasks. A one-way repeated-measures ANOVA (N=7 mice – N.B. only 7 of the 11 mice were presented with all 40 tasks) showed a main effect of Task F=3.10 P=0.0100, df1=7, df2=42. f) Animals readily performed zero-shot inference on the first trial of late tasks but not in early tasks. The proportion of tasks in which animals took the most direct path from ***d*** to ***a*** on the very first trial is compared to the same measure but for premature returns from ***c*** to ***a*** and ***b*** to ***a*.** Wilcoxon test; Early tasks: N=11 mice, statistic=15.5, P=0.407; Late tasks: N=11 mice, statistic=2.0, P=0.009 All error bars represent the standard error of the mean

We first asked whether animals learned optimal sequences leading to individual goals. To assess performance, we quantified either the ratio of the distance taken between two goal locations compared to the shortest possible distance (“Relative path distance”), or the proportion of trials where one of the shortest routes was taken (“Proportion shortest”). On individual tasks, mice converged on a near-optimal policy that routinely took them between goal locations via close-to-shortest routes (Figure 1c). This converged policy was highly stereotyped, with animals taking only a subset of the available shortest routes (Extended data fig 1b). In the first 10 tasks, we allowed animals to perform as many trials as needed to converge on peak performance (Figure 1c; 70% shortest path transitions or 200 trial plateau; mean number of trials per task: 325 ±16). Subsequently, to encourage generalisation of task structure, and to allow us to record from multiple tasks in a day, we moved animals to a high task regime (tasks 11-40). Animals experienced 3 new tasks per day and hence could only complete a fraction of the trials needed to reach peak performance (mean number of trials per task: 38 ± 3). Despite this, animals still performed markedly above chance (Figure 1d). Suboptimal performance was associated with a persisting preference for taking routes around the maze for which animals had an *a priori* bias to take before exposure to the task (Extended data figure 1c). Nevertheless, animals’ performance improved across even the early trials of a given task (Figure 1d).

We next asked whether mice learn the higher order, abstract task structure. Initial performance on the earliest trials improved with the number of tasks completed (Figure 1d; Extended data figures 1d,e). This improvement was seen even on the very *first* trial of each new task (Figure 1e). This effect is consistent with animals transferring knowledge of task structure to rapidly learn new sequences. Notably, this task allows for a *direct* test of this structural knowledge transfer because of the ABCD loop. After discovering the 4 reward locations (***a*,*b*,*c*** and ***d***) on the first trial of a new task, animals that understand the task structure should then return directly to ***a***. On the first trial, this transition (***d* → *a***) has never been experienced, so cannot be executed through repetitive learning or memory. Instead it must reflect a zero-shot inference based on abstract knowledge of the ABCD task structure. Remarkably, we found that experienced animals took the shortest path between ***d*** and ***a*** on the first trial more often than chance and more readily than premature returns to ***a*** from ***b*** or ***c*** (Figure 1f). This was not explained by any pre-existing biases in the animals’ exploration of the maze (Extended data fig 1f), differences in analytical chance levels (Extended data fig 1g) or differences in the distances of the ***d***-to-***a*** transition compared to those for ***c***-to-***a*** and ***b***-to-***a*** (Extended data fig 1h). Moreover, this zero-shot inference was associated with animals returning to the start of the loop after 4 rewards (***d***-to-***a***) rather than to later points of the loop (***d***-to-***b*** or ***d***-to-***c*;** Extended data fig 1i). Thus animals not only waited until 4 rewards were obtained before returning to ***a*** (Figure 1f) but also more readily returned to ***a*** than to other reward locations after 4 rewards (Extended data fig 1i). Overall these findings suggest that mice learn an abstract, task-defined behavioural structure nesting multiple goals.

### Progress to goal is a primary feature of Frontal task structure representations

Animals learn the sequences leading to individual goals, and the abstract structure organising these goals. What are the basic neural underpinnings of this hierarchical learning? In order to perform such complex sequences, animals must track their position not only in the physical space they are navigating, but crucially also their “progress” in task space, that is the stage the animal has reached in a sequence of goal-directed behaviours. Are mFC neurons tuned to task progress?

We used silicon probes to record the activity of mFC (prelimbic) neurons (Figure 2a; Extended data figure 2a) from animals (N=5) performing late tasks (tasks 21-40), a stage where we see robust evidence for task structure knowledge (Figure 1d-f). Each recording day comprised 3 sessions with 3 new tasks (X, Y and Z) and a subsequent fourth session in which the first task was repeated (X’). In addition, we recorded sleep sessions at the start and end of the day and in between every session. We used a generalised linear model to tease out the variables explaining mFC neuronal activity. The large majority (80%) of mFC neurons were strongly and consistently tuned to the animal’s progress towards a rewarded goal, regardless of where the reward was spatially (Figure 2b-f). Neurons tiled goal-progress space: some fired immediately after the animal reached its goal (early goal-progress cells), others were most active between two goals (intermediate goal-progress cells) and others still just before a goal was reached (late goal-progress cells; Figure 2c-e). This goal-progress tuning was highly invariant across tasks with distinct reward locations (Figure 2e) and not explained by simple monotonic tuning to speed nor acceleration (Figure 2f). Furthermore, these neurons fired in relation to goal-progress relative to goals independently of elapsed time or physical distance (Figure 2f). Thus, mFC neurons exhibit stimulus-invariant tracking of progress towards individual goals.

**Figure 2 –.**
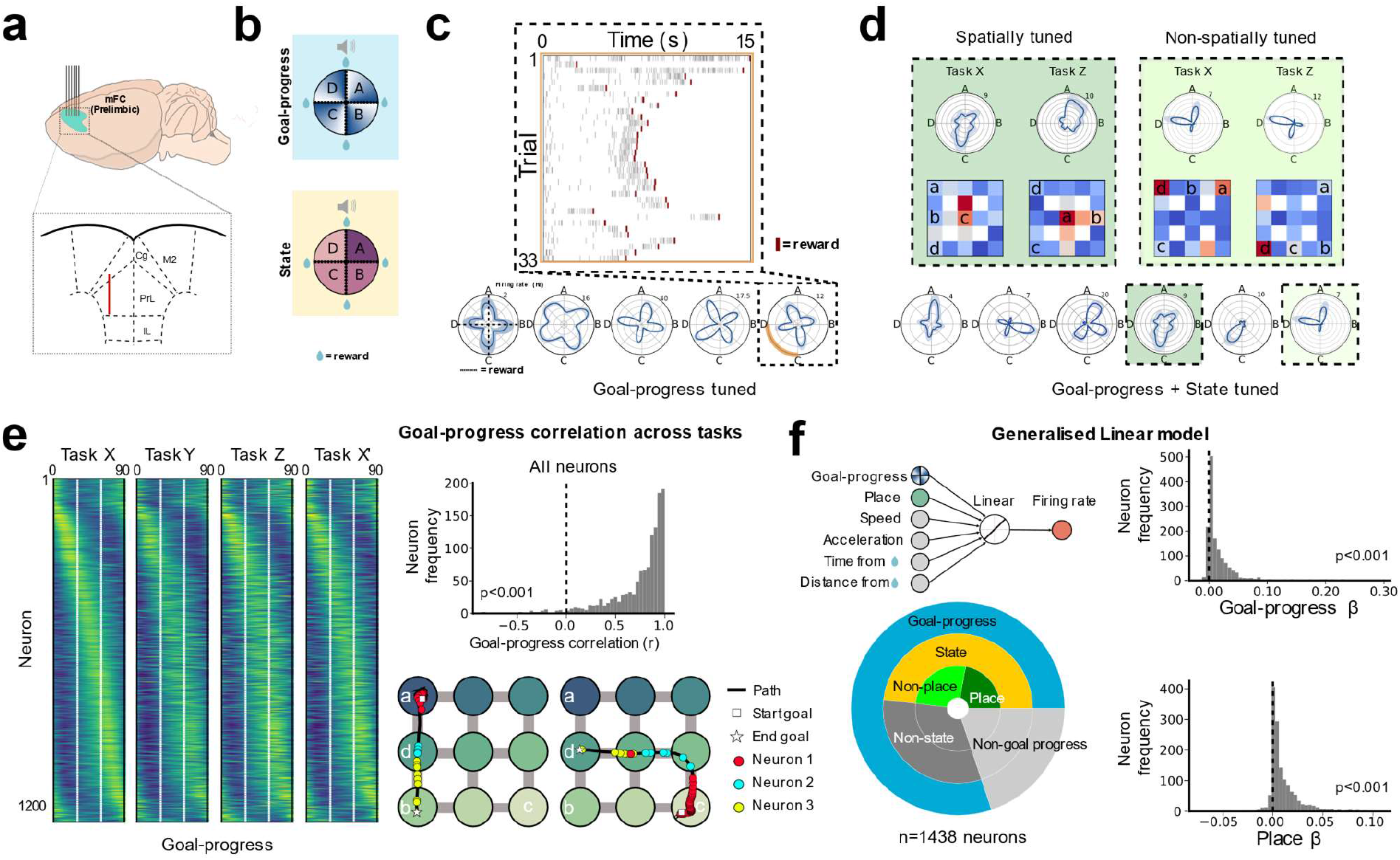
Progress-to-goal is a key feature of task-tuned neurons in the medial Frontal Cortex. a) Multi-unit recording set-up: animals were implanted with silicon probes targeting 1mm of the anterior-posterior extent of the prelimbic region of the medial Frontal cortex. Inset: schematic of a coronal slice through the mFC showing electrode placement in the prelimbic region. b) Schematics of polar plots showing a projection of neuronal activity onto the circular task structure. “Goal-progress” refers to how much progress an animal has made towards a rewarded goal location as a percentage of the time taken to reach this location, while “State” refers to progress in the overall ABCD loop comprising all 4 goals. The radial axis represents a neuron’s firing rate, while the angular axis represents progression along the task states. Dashed lines along the cardinal directions represent the times reward was obtained (goal was reached) in each state, with the vertical line at zero representing reward ***a***, i.e. the start of state A; going clockwise, the remaining dashed lines represent reward locations ***b***, ***c*** and ***d***, and hence the starts of states B C and D respectively. To represent task position, bins along the angular axis increment with relative progress between goals rather than raw elapsed time. c) Neurons display consistent tuning to the progress of the animal’s relative to any goal (“goal-progress tuned”). Inset: a raster plot of firing activity in one state “C” of a goal-progress cell, showing firing consistently shortly before a goal is reached. d) Some goal-progress-tuned cells are additionally modulated by the state in a given task (“goal-progress + State tuned”). Inset: Polar plots and spatial maps for two goal-progress + State tuned neurons across two distinct task configurations. The neuron on the left is spatially tuned while the neuron on the right is non-spatially tuned. e) Goal-progress tuning is consistent across tasks that differ in reward locations. Left: The average firing rate vector of all neurons relative to an individual goal (from goal “n” to goal “n+1”; averaged across all states). Animals experience 3 tasks per day (tasks X, Y and Z) and then a further 4th session which task X is repeated (task X’). Each row represents a single neuron and the neurons are arranged on the y axis by their peak firing goal-progress in task X. This alignment is largely maintained in tasks Y and Z as well as a later session of the first task (X’). White dashes indicate early intermediate and late goal-progress-cutoffs. Top right: A histogram showing the mean goal-progress-vector correlation across tasks for each neuron. One-sample T test against 0: N=1230 neurons; statistic=91.56; P=0.0, df=1229. Note that the neurons used in this panel are those that were tracked for all 4 sessions on a given day. Bottom right: two example paths, each between a pair of rewarded goals and overlaid spiking of 3 goal-progress-tuned mFC neurons tuned to early goal-progress (neuron 1) intermediate goal-progress goal-progress (neuron 2) and late goal-progress (neuron 3) regardless of the goal locations or trajectory taken. f) Top left: a schematic of the variables inputted into a generalised linear model that predicts neuronal activity across tasks and states. Bottom left: Pie-chart showing the results of a generalised linear model capturing variance as a function of goal-progress, place, speed, acceleration, time from reward and distance from reward. Plot shows proportions of neurons with significant regression coefficient values for goal-progress, place and their conjunctions. It also shows proportions of state tuned neurons derived from a separate z-scoring analysis (More details in Methods under “Tuning to basic task variables”). Proportion of all neurons that are goal-progress cells: 80%; Two proportions test: N=1438 neurons, z=40.7, P<0.00. Proportion of all neurons that had state tuning in at least one task: 56% Two proportions test: N=1438 neurons, z=29.5, P<0.001. 87% of all state-tuned neurons were also goal-progress tuned; Two proportions test: N=798 neurons, z=32.8, P<0.001. Proportion of goal-progress-tuned neurons that also showed significant state preference in at least a single task: 60%; Two proportions test: N=1152 neurons, z=28.2, P<0.001. Proportion of neurons tuned to goal-progress and state that were also tuned to place: 46%, Two proportions test: N=692 neurons, z=17.4, P<0.001. Top right: A histogram showing the mean regression coefficient values for *goal-progress* as a regressor across task/state combinations for each neuron. One-sample T test against 0: N=1438 neurons; statistic=20.52; P=2.71×10^−82^, df=1437. Bottom right: A histogram showing the mean regression coefficient values for *place* as a regressor across task/state combinations for each neuron. One-sample T test against 0: N=1438 neurons; statistic=24.55; P=1.96×10^−111^, df=1437

The invariant goal-progress tuning of mFC neurons is consistent with this region tracking progress in relation to *individual* behavioural goals, rather than physical distance or time. However, such goal-progress tuning alone is insufficient for tracking position in the higher order task structure organising multiple goals. We therefore leveraged the hierarchical structure of our ABCD task to ask whether mFC neurons are tuned to a given *state* in an individual ABCD task (e.g. state B). We found such “state” tuned neurons in abundance (56% of all neurons had state tuning in at least one task; Figure 2d,f; Extended data figure 2b). Intriguingly, the large majority (87%) of these state tuned neurons were also goal-progress tuned (Figure 2f). Thus, task-state-tuned neurons are largely a subset of the more prevalent goal-progress-tuned neurons. A corollary of this is that a large proportion (60%) of goal-progress-tuned neurons also showed significant state preference in at least a single task (Figure 2d,f; Extended data figure 2b). Moreover, 46% of neurons tuned to both goal-progress and state were additionally tuned to place (Figure 2d,f; Extended data figure 2b). Thus, for 54% of cells with state and goal-progress tuning, state-preference was not explained by tuning to the animal’s current spatial location (Figure 2d,f; Extended data figure 2b). These findings suggest that progress-to-goal is a key determinant of mFC neuronal firing and that a subset of these goal-progress cells are additionally tuned to a given state in a given task.

### Modular organisation of mFC task structure mapping

Neurons in the mFC delineate task state on individual tasks. One view holds that, in order to support transfer of task structure knowledge across distinct examples, such state tuning should explicitly generalise across different tasks^32,34^. This would then allow an associative solution to the abstraction problem, whereby “abstract state” cells would (re)bind flexibly to neurons representing specific spatial goals in order to encode new tasks^32,34^. To investigate this possibility, we asked whether the state tuning of mFC neurons, which reflects the animal’s position in the ABCD loop of one task, is invariant across tasks. Intriguingly, rather than invariant state tuning, we found that neurons consistently remapped in task space. In contrast to their goal-progress tuning, which was invariant across tasks (Figure 2), mFC neurons did not retain their state preference across tasks (Figure 3a,b). Notably, state-tuned neurons with no discernable spatial tuning also remapped across tasks (Extended data Figure 3b). Crucially, when we compared state tuning across different sessions of the same task (X and X’), state tuning was highly conserved, despite two intervening sessions with different tasks and different state tuning (Figure 3b). This was also seen when only considering non-spatial neurons (Extended data Figure 3b). Overall these results indicate that, while neurons in the mFC invariantly map progress to a given goal, they do not invariantly map abstract task states in a higher order structure relating the sequential goals.

**Figure 3 –.**
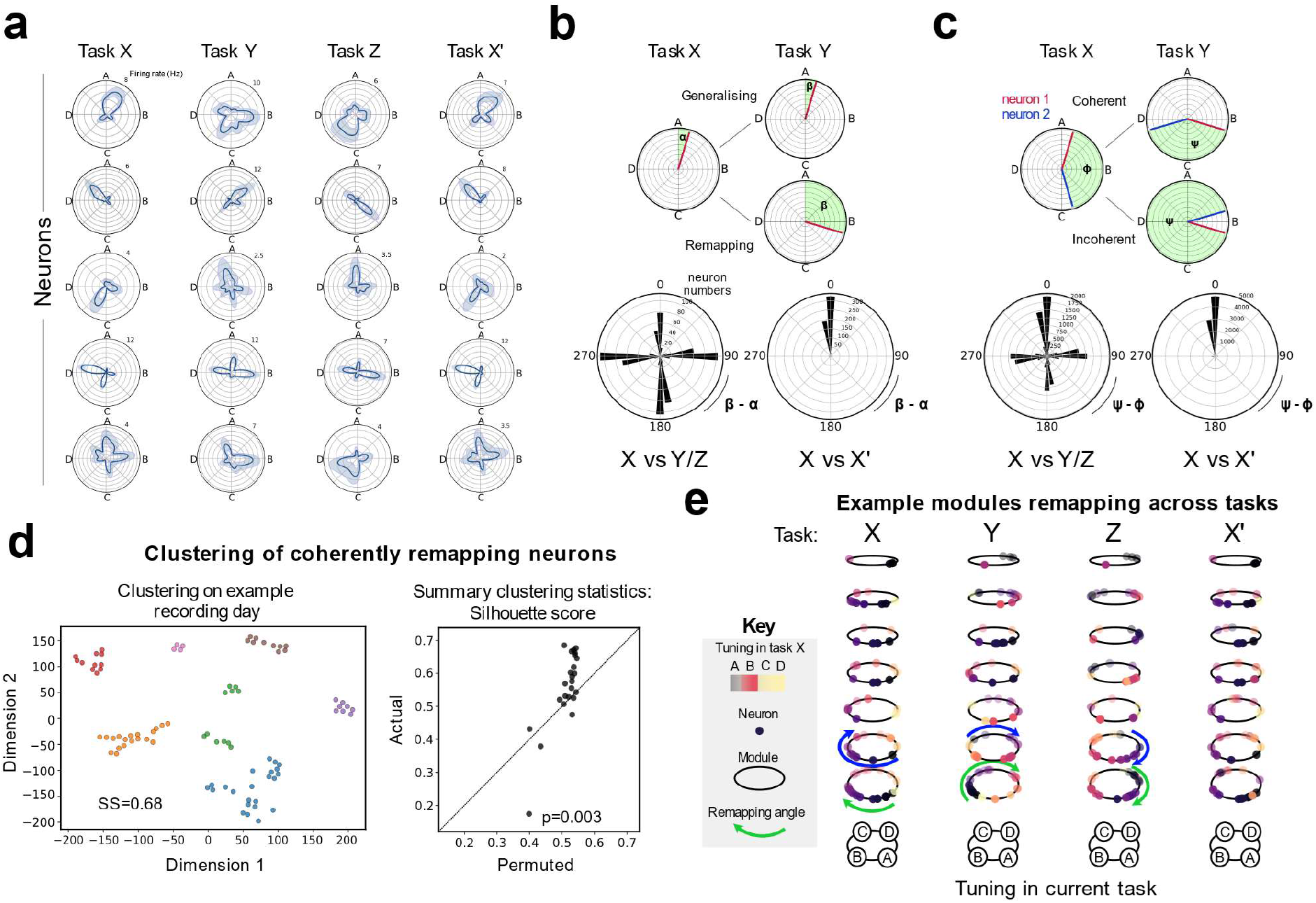
Medial Frontal neurons are organised into task-space modules. a) Polar plots showing state tuning of example neurons across multiple tasks. Each row is a neuron and each column is a session. Neurons readily remap their state tuning but maintain their goal-progress preference across tasks. State preference is maintained across different sessions of the same task (X vs X’). b) Top: A schematic showing how the difference in tuning angles for the same neuron across sessions is quantified. Bottom left: Polar histograms show that state-tuned neurons remap by angles close to multiples of 90 degrees, as a result of conserved goal-progress tuning and the 4 reward structure of the task. No clear peak at zero is seen relative to the other cardinal directions when comparing sessions spanning separate tasks (Two proportions test against a chance level of 25% N=831 neurons; mean proportion of generalising neurons across one comparison (mean of X vs Y and X vs Z)=20%, z=9.04, P<0.001: i.e. significantly lower than chance). Bottom right: Neurons maintain their state preference across different sessions of the same task (X vs X’ Two proportions test against a chance level of 25% N=624 neurons; proportion generalising=87%, z=29.0, P<0.001). c) Top: A schematic showing how the difference in relative angles between pairs of neurons across sessions is quantified. Bottom left: Polar histograms show that the proportion of coherent pairs of state-tuned neurons (comprising the peak at zero) is higher than chance but less than 100%, indicating that the whole population does not rotate coherently. Two proportions test against a chance level of 25% N=14930 pairs; mean proportion of coherent neurons across one comparison (mean of X vs Y and X vs Z)=32%, z=59.8, P<0.001). Bottom right: As expected from panel b, the large majority of state-tuned neurons keep their relative angles across sessions of the same task (X vs X’; Two proportions test against a chance level of 25% N=10722 pairs; proportion coherent=79%, z=109.2, P<0.001). d) Left: Example from a single recording day showing the result of tSNE embedding and hierarchical clustering derived from a distance matrix quantifying cross-task coherence relationships between state-tuned neurons. Each dot represents a neuron. Right: Summary silhouette scores for the clustering for real data compared to permuted data that maintains the neuron’s goal-progress preference and initial state distribution. Each dot is a recording day. Wilcoxon test: N=24 recording days; statistic=50.0, P=0.003 e) Visualisation of tuning relationships between clusters computed in a single recording day. Each dot is a neuron and each ring is a cluster derived from the analysis in panel d. The colour code represents the tuning of the neurons in task X. The x,y position defines the tuning in each task. The z position corresponds to cluster ID. Note that the ordering along the z axis is arbitrary. Neurons rotate (remap) in task space while maintaining their within-cluster tuning relationships but not cross-cluster relationships across tasks.

While remapping precludes a model in which neurons are invariantly tuned to the ordinal position of each goal, it may nevertheless still reflect an invariant structure. Neurons could maintain a consistent ring-like arrangement which maps the abstract task structure but rotates coherently across different tasks (e.g. if all A neurons become C neurons then all D neurons should become B neurons…etc). Such a representation could still allow tracking abstract task position and the retrieval of concrete behavioural states through an associative mechanism. We therefore asked whether the remapping of mFC neurons across tasks reflects a coherent structure between neurons in the population. We measured the degree of coherence between pairs of mFC state-tuned neurons: how likely is it that a pair of neurons will maintain their relative state preference across tasks? Only a small proportion of pairs, but significantly above chance level, showed coherent remapping across tasks (Figure 3c). This pairwise coherence was true for non-spatial neurons (Extended data figure 3c) and when neurons were tuned to distant points in task space (Extended data figure 3d). Moreover, coherent pairs of neurons showed a trend towards being closer anatomically (Extended data figure 3e). The partial pairwise coherence between mFC neurons suggests that such neurons might be organised into modules; groups of neurons that maintain their tuning relationships across tasks, akin to the modular arrangement of grid cells mapping physical space^36^. To investigate this, we used a clustering approach. We defined a distance metric between pairs of cells which assigned low distances between cells that remapped coherently between tasks, and high distance between cells that remapped incoherently. We then applied a low dimensional embedding (tSNE) on the resulting distance matrix, followed by hierarchical clustering. mFC neurons were significantly clustered (Figure 3d,e), indicating that they were organised into modules which remap coherently across tasks. Overall, these findings suggest that mFC neurons don’t generalise their state tuning relationships as a coherent whole but are instead organised into modules that conserve their within-module, but not between-module, tuning relationships across tasks.

### Structured memory buffers: a unified model for behavioural schema and sequence memory

The modular arrangement of mFC neurons indicates that the entire population isn’t anchored to a single invariant reference point (e.g. state A). Instead, because they rotate independently, each module is anchored to a distinct reference point. What could these reference points be? We reasoned that the strong tuning of some frontal neurons to both goal-progress and spatial location (Figure 2) could offer an answer. Each module of neurons could be anchored to a particular conjunction of goal-progress and place (e.g. early goal-progress at location 1; Figure 4a) through a subset of “*anchor*” neurons tuned to that specific goal-progress/place combination. The other “*non-anchor*” neurons in the module would then each fire at a specific lag in task-space from when the animal visits the goal-progress/place anchor (Figure 4a). This amounts to a change in reference point for each module: from the abstract states (ABCD) to a module-specific anchor. Under this scheme, the apparent remapping seen when aligning activity to abstract states (ABCD; Figure 3) occurs because animals visit the goal-progress/places in a different sequence in each task (Figure 4b,c). If in one task location 1 is rewarded in state A and in another the same location is rewarded in state C, a module anchored to early goal-progress in location 1 would appear to rotate by 180 degrees across tasks when aligned to the abstract states (Figure 4c). All neurons on this module, not just the anchor neurons, would rotate by the same amount. The anchor neurons, which are tuned to a particular goal-progress/place conjunction, would remap in a way explained by their spatial tuning. However, neurons further from the anchor along a given module would remap in one task in a way that is not explained by their spatial map in another task (Figure 4 b,c) just as seen empirically (Extended data figure 3b,c). Each module with a different anchor would rotate by a different amount depending on when the goal-progress/place encoded by the anchors are encountered in the behavioural trajectory of the current task. Thus, we posit that the true invariance of state neuron activity can only be seen when realigning their activity to their putative goal-progress/place anchors (Figure 4a).

**Figure 4 –.**
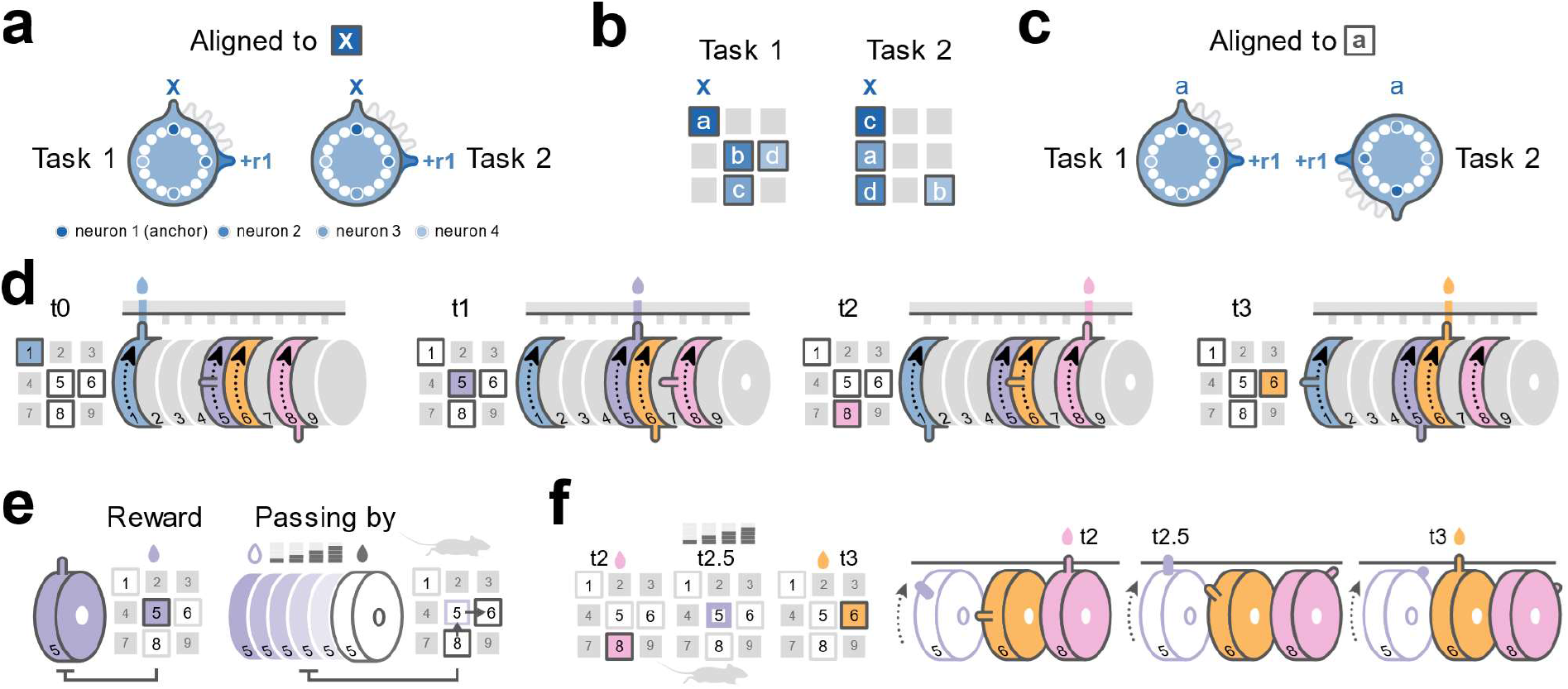
The Structured Memory Buffers model. a) A hypothetical ring-shaped module of neurons “anchored” to location 1 on the maze (marked by an X) in early goal-progress (i.e. during reward consumption). The ring therefore represents a memory buffer for this anchor. Aligning the ring of neurons by this anchor reveals the invariant relationships between neurons across any two tasks where location 1 is rewarded. 4 neurons are highlighted; the dark blue anchor neuron (neuron 1) and 3 other neurons firing at lags of 90 degrees (i.e. one reward away (one state away); neuron 2), 180 degrees (i.e. two rewards away; neuron 3) and 270 degrees (i.e. three rewards away; neuron 4) from the anchor. Note that other neurons at all other task lags, not just multiples of 90 degrees, are also present on the ring (white circles). A bump of activity is initiated when the goal-progress/place anchor is visited (top) and moves around the ring paced by the animal’s progress in task space. In this task, because there are four rewards, the bump moves by a total of 90 degrees around the ring after each reward (e.g. the bump reaches neuron 2 after the animal has received 1 further reward since receiving reward in location 1). b) Two example ABCD tasks that share one reward location (location 1, marked by an X). The shaded regions correspond to the hypothetical spatial firing fields of each of the 4 neurons shown in panel a - shading corresponds to the shading of the neurons on the ring in panel a. While the anchor neuron (neuron1; dark blue) fires consistently at the same goal-progress/place conjunction across tasks, other neurons (neurons 2-4; lighter shades of blue) will fire in different locations in the two tasks. This is because they primarily encode elapsed task progress from early reward in location X, rather than physical space. So for example, neuron 4 (the lightest shade of blue) consistently fires 3 rewards from the time the animal got reward in location 1: this means it fires in location 6 (reward ***d***) in task 1 as it was 3 rewards from reward in location 1 (reward ***a***); in task 2 this neuron fires in location 9 (reward ***b***) as this is 3 rewards from reward in in location 1 (reward ***c***). c) The same ring, when aligned by the abstract task states (e.g. aligned to start of state A; i.e. reward in location ***a***), appears to rotate by 180 degrees across tasks. This is a direct result of reward in location X corresponding to the start of a different state in each task (state A in task 1 and state C in task 2). d) A time series showing the flow of activity along 4 rings, each anchored to one of the 4 rewarded locations in task 1. A bump of activity is initiated when the goal-progress/place anchor is visited (top) and moves around the ring paced by the animal’s progress in task space. When it circles back close to the start, it biases the animal to return to this anchor. Multiple rings have active bumps at any one time, thereby simultaneously tracking the history of different goal-progress/place anchor visits (i.e. the history of previous behavioural steps). e) As well as rings tracking task-progress from behavioural steps involving a rewarded place (a conjunction of a place with early goal-progress), there are also rings tracking task-progress from places conjoined with intermediate and late goal-progress. The anchors of these rings are activated when the animal passes through a location, not when it is rewarded, but at a defined, non-zero progress relative to the goal. d) Non-zero goal-progress anchored rings (e.g. purple outline) allow tracking task-progress from behavioural steps in between two goals. Hence, across all rings, a history of the entire sequence of steps taken by the animal, not just the sequence of reward locations, is encoded at any one point in time.

A key implication of this model is that each module is a *memory buffer* for visits to a particular anchor. Because it is a combination of location and goal-progress, an anchor represents a particular behavioural step. The flow of activity along a given module answers the following question: how much of the overall *task* (i.e. the 4 reward sequence in our ABCD task) has elapsed since visiting a given behavioural step? (for example since visiting early goal-progress in location 1; Figure 4a,d) Because there are neurons for every task-space lag from a given behavioural step within each module, and modules for every possible behavioural step, there are always a subset of active neurons that are tied to each visited behavioural step (Figure 4d). The instantaneous mFC activity therefore always encodes the entire sequence of behavioural steps. Moreover, because there are anchors representing behavioural steps at intermediate goal-progress, not only at rewards, this encoded sequence is the entire behavioural sequence executed by the animal (i.e. the route taken through the maze), not just the sequence of 4 reward locations (Figure 4e,f). These modules are organised by task structure in two ways. Firstly, the strong goal-progress tuning of mFC neurons means that activity on the module evolves as a function of the number of goals obtained (i.e. the number of goal-progress cycles completed). The modules therefore track true task-progress rather than other dimensions such as elapsed time or distance travelled. Secondly, a module is shaped by the structure of the task, in our case a four reward ring. Once an animal completes a full trial (i.e. traverses 4 goals in the ABCD task) since it last visited a given behavioural step, activity along the module will circle back to the anchor point (Figure 4d,f). This means that the memory buffers are “structured”: they are internally organised to reflect the abstract structure of the task, a 4-reward loop in our case. We refer to this over-arching mechanism as the <u>S</u>tructured <u>M</u>emory <u>B</u>uffer (SMB) model.

The SMB model posits that activity along mFC modules could be used to guide the execution of task-paced behavioural sequences. Once an animal completes a full trial since it last visited a given behavioural step, either the anchor neurons themselves or neurons close to them along a given module could be used as output neurons that bias the animal to return to the behavioural step represented by the anchor. Collectively, activity dynamics along *all* active modules would allow retrieval of a sequence of behavioural steps (policy) to solve a given task. Consequently, any new sequence with the same structure can be mapped in a programmable way, simply by reconfiguring the order in which modules are activated. These network dynamics are sufficient to encode new tasks, without needing new plasticity. For simplicity, we have assumed so far that each neuron has a single anchor with a single lag. However, in principle the same computational logic can be used even when individual neurons respond to a combination of different anchors and lags. The read-out in such a scenario would involve combinatorial activity across anchor neurons across multiple modules.

The SMB model is computationally attractive because it unifies frontal cortex functions in behavioural schema formation and sequence memory, while offering a programmable way of encoding and retrieving sequential policies. It is also empirically attractive because it explains the spatial-tuning-independent state preference (Figure 2d,f), remapping (Figure 3a,b) and the modular arrangement (Figure 3c-e) of the neurons across tasks. Crucially, the model makes a number of new empirical predictions about the tuning of single neurons, their relationship to behavioural choices and their organisation at the population level. We explore these predictions in turn below.

#### 1- Mnemonic fields in task space

The SMB model proposes that, instead of neurons being anchored to the abstract task states (A,B,C or D), they instead encode how much task-space has elapsed (i.e. how much task progress has been made) since the animal visited a specific behavioural step. These neurons should therefore maintain an invariant task-space lag from their module’s anchor (behavioural step) across tasks. For example, a neuron would always fire 2½ states from the time the animal received reward (i.e. early goal-progress) at location 3, regardless of where the animal is physically at this point. We tested this prediction using three complementary approaches. In all of these analyses, we concatenated two recording days, giving a total of up to 6 new tasks per neuron.

In the first approach, we implemented a linear regression model to predict the state tuning of neurons across tasks. For each neuron, the model describes state-tuning activity as a function of all possible behavioural steps (conjunctions of goal-progress and place) and task lags from each possible behavioural step. Thus a neuron could fire at a particular behavioural step but also at a non-zero lag in task space from this behavioural step. We trained the model on all tasks but one (training tasks) and then used the resultant regression coefficients to predict the activity of the neuron in a left out (test) task. To ensure our results are due to task-lag preference and not the powerful effect of goal-progress tuning (which is easy to predict across tasks due to its invariance; Figure 2 e,f), both training and cross-validation were only done in the preferred goal-progress of each neuron. Using this approach we were able to predict the state preference of most state-tuned neurons on individual tasks as evidenced by a strong rightward (positive) shift in the correlation between predicted and actual state tuning in the test task (Figure 5a,b; Extended data Figure 4a; Extended data Figure 5a). Crucially, this rightward shift was also seen when only considering activity at non-zero lags relative to all anchors (Figure 5b, Extended data Figure 5a,b). Neurons were distributed across all lags from the anchors, with an overrepresentation of zero lag neurons, corresponding to the anchors (behavioural steps) themselves (Extended data Figure 5c). Moreover, we found neurons anchored to all goal-progress/place combinations (Extended data Figure 5d).

**Figure 5 –.**
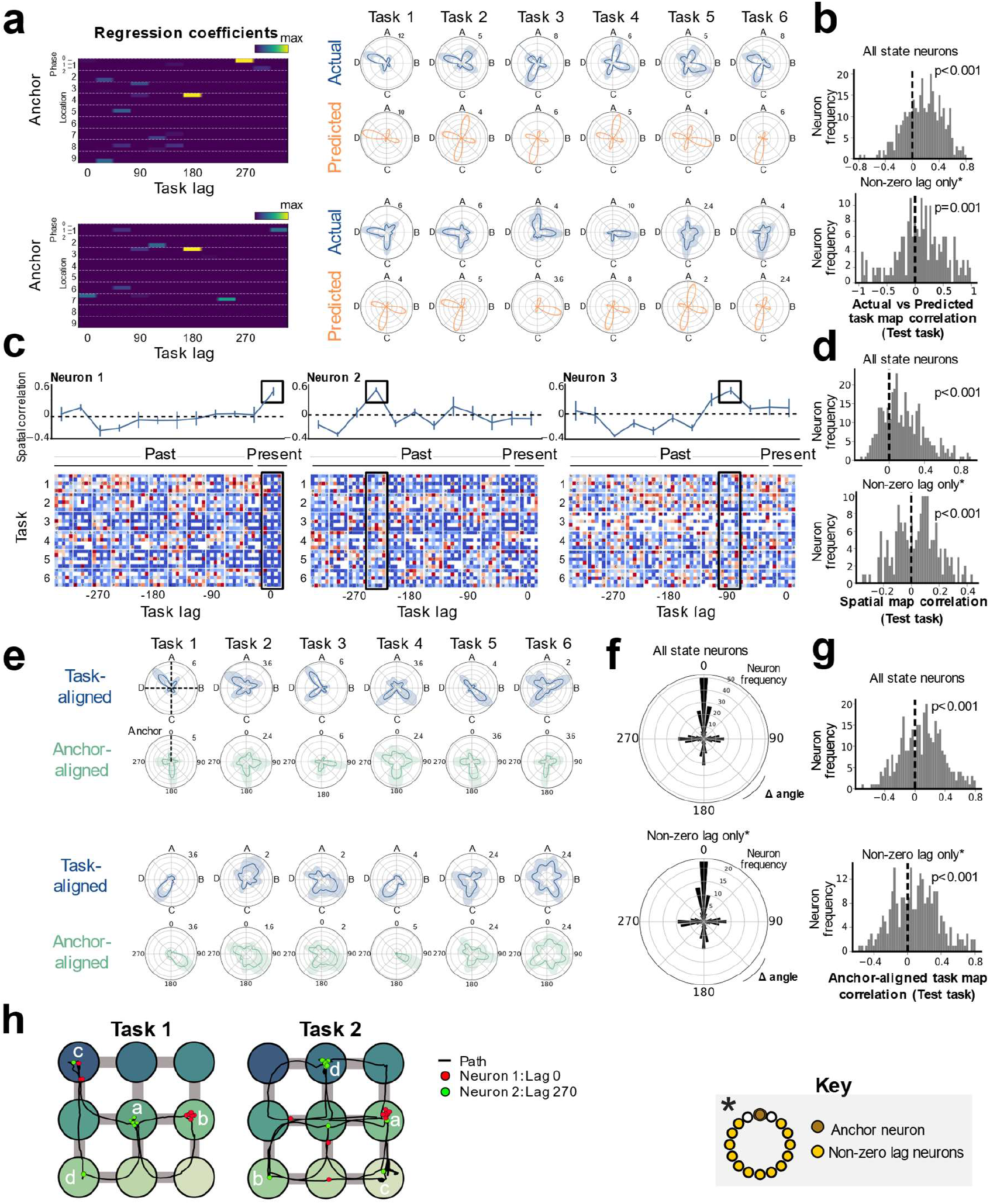
Medial Frontal neurons track task-progress from specific behavioural steps. a) Regression analysis reveals neurons with mnemonic fields in task space from a given goal-progress/place conjunction, alongside neurons directly tuned to a goal-progress/place. This allows predicting state-tuning and its remapping across tasks, The regression coefficients are shown on the left with the actual and predicted activity of the neurons shown on the right. Left: The top neuron has its highest coefficients at 180 degree task lag from early goal-progress in location 4 and 270 degrees task lag from early goal-progress in location 1.The bottom neuron has its highest coefficients at 180 degree task lag from early goal-progress in location 3 and 240 degrees task lag from intermediate goal-progress in location 7. Right: Actual (blue) and model-predicted (orange) activity across 6 tasks b) Histograms showing the right shifted distribution of mean cross-validated correlation values between model-predicted (from training tasks) and actual (from a left out <u>test task</u>) activity. Top: this correlation is shown for all state-tuned neurons and Bottom: only state-tuned neurons with non-zero-lag firing from their anchors (i.e. state-tuned neurons with all of the three highest regression coefficient values at non-zero lag (30 degrees either side of the anchor). To avoid contamination due to potential residual spatial-tuning, for non-zero lag neurons, we only use regression coefficient values more than 30 degrees in task space either side of the anchor point to predict the state tuning of the cells. T test against 0: All state-tuned neurons N=340 neurons, statistic=11.1, P=1.36×10^−24^, df=339; Non-zero lag state-tuned neurons N=194 neurons, statistic=3.26, P=0.001, df=193. c) Lagged spatial field analysis. Example plots showing spatial maps for 3 neurons. Bottom: Each row represents a different task and each column a different lag in task space. The activity of each neuron is plotted as a function of the animal’s current location (far right column for each cell) and at successive task space lags in the past for the remaining columns. For example, the second to last column is the firing of the cell in relation to where the animal was 30 degrees (1/3rd of a state) back from the current point in task space, 7th to last column is the firing of the cell in relation to where the animal was 180 degrees (2 states) back from the current point in task space…etc. Top: the correlation of spatial maps across tasks at each lag. The neuron on the left is an anchor cell (goal-progress/place cell), as seen by the correlation peak at zero lag, while the middle and right-most neurons are neurons lagged by 240 and 90 degrees from their anchors respectively. d) Histograms showing the right shifted distribution of the mean cross-validated spatial correlations between maps at the preferred lag (from training tasks) and the spatial map at this lag from a left out <u>test task</u> for all state-tuned neurons (top) and only non-zero lag neurons (bottom). Non-zero lag neurons are those with a peak spatial map correlation more than 30 degrees in task space either side of zero lag in the training tasks. T test against 0: All state-tuned neurons N=350 neurons, statistic=13.0, P=1.24×10^−31^, df=349; Non-zero lag state-tuned neurons N=179 neurons, statistic=3.48, P=6.39×10^−4^, df=178. e) Single anchor alignment analysis. Top (blue) plots for each neuron shows activity aligned by the abstract states (with the dashed vertical line at zero representing reward *a*, i.e. the start of state A; going clockwise, the remaining dashed lines represent reward locations ***b***, ***c*** and ***d***, and hence the starts of states B, C and D respectively). Neurons appear to remap in task space across tasks. Bottom (green) plots for each cell show that it is possible to find a goal-progress/place anchor that consistently aligns neurons across tasks (the zero dashed vertical line corresponds to visits to the goal-progress/place anchor). The top Neuron aligns best to its anchor close at 180 degrees while bottom neuron aligns best to its anchor at 120 degrees. f) Polar histograms showing the cross-validated alignment of neurons by their preferred goal-progress/place anchor (calculated from training tasks) in a left-out test task. The top plot is for all state-tuned neurons while the bottom plot only includes non-zero lag neurons. Non-zero lag neurons are those with a peak firing rate more than 30 degrees in task space either side of the anchor. Two proportions test against chance (25%): All state-tuned neurons: Proportion generalising: 38.1% N=350 neurons, z=10.66 P<0.001; Non-zero lag state-tuned neurons: Proportion generalising: 38.5% N=269 neurons, z=9.43, P<0.001. g) Histograms showing the right shifted distribution of the mean cross-validated task map correlations between state-tuned neurons aligned to their preferred goal-progress/place anchor (from training tasks) and the task map aligned to this same goal-progress/place anchor from a left out <u>test task.</u>. This is shown for all state-tuned neurons (top) and only non-zero lag state-tuned neurons (bottom). Non-zero lag neurons are those with a peak firing rate more than 30 degrees in task space either side of the anchor. P values are from T tests relative to 0. T test against 0: All state-tuned neurons N=350 neurons, statistic=6.12, P=2.53×10^−9^, df=349; Non-zero lag state-tuned neurons N=269 neurons, statistic=5.44, P=1.89×10^−7^, df=268. h) Two example paths each spanning an entire trial across two distinct tasks and overlaid spiking of 2 mFC state-tuned neurons anchored to early goal-progress in location 6 (middle right location on the maze). Neuron 1 is an anchor neuron and is hence tuned to this goal-progress/place. Neuron 2 fires with a lag of roughly 270 degrees in task space from the anchor and so fires 3 states after the animal visits early goal-progress in location 6. Thus neuron 2 fires when animals get reward in location 5 in task 1 (reward ***a***) as this is 270 degrees (3 states) in task space from reward in location 6 (reward ***b***). In task 2, this neuron fires most in location 2 (reward ***d***) which is also 270 degrees (3 states) from reward in location 6 (reward ***a***). For visualisation purposes, spikes were jittered to ensure directly overlapping spikes are distinguishable. All error bars represent the standard error of the mean

The second approach involves assessing the spatial tuning of mFC state-tuned neurons. Classically, spatial maps relate the activity of neurons to where the animal is located in the *present*, that is with zero lag in task space^1^. Our model proposes the existence of neurons tuned to where the animal was a set amount of task lag in the *past*. The logic of this is that, while spatial neurons should consistently fire at the same locations(s) in the present (i.e. at zero lag from the present), neurons that track a memory of the anchor will instead consistently fire in relation to when the animal visited the anchor at a fixed lag in task space. They will therefore have a peak in their cross-task spatial correlation at a fixed, non-zero task lag in the past (Figure 5c; Extended data Figure 4b). Unlike the first analysis, this approach assumes a single lag from an anchor per neuron, but still allows the anchor to be a combination of locations. To quantify this effect, we again used a cross-validation approach, this time using training tasks to calculate the lag at which cross-task spatial correlation was maximal, and then measuring the correlation between the spatial maps in the left out (test) task and the training tasks at this lag. This correlation was again strongly right-shifted (Figure 5d), which was the case even when considering only neurons with non-zero lag relative to the animal’s current location (Figure 5d, Extended data Figure 5e,f).

The third approach is effectively the reverse of the second. Instead of comparing spatial tuning aligned to particular lags, it compares lag-tuning when aligned to particular anchor visits (Figure 5e-h). Unlike the first two analyses, this approach assumes a single anchor per neuron. We fitted the anchor by choosing for each neuron the goal-progress/place conjunction which maximises the correlation between lag-tuning-curves in the training tasks, and again used cross-validation by assessing whether this anchor leads to the same lag-tuning in a left out task (Figure 5e; Extended data Figure 4c). Using this cross-validation approach, we found significant alignment between firing distances from the best anchor in the training and test tasks (Figure 5f), which was crucially seen even when only considering non-zero lag neurons (Figure 5f). This was in stark contrast to aligning the activity by abstract state (ABCD), which showed no peak at zero relative to other cardinal directions (Figure 3b). To quantify the degree of alignment further, we measured the correlation between the activity of neurons in their preferred goal-progress between the test and training tasks. The resultant distribution was again right-shifted, even for non-zero lag neurons (Figure 5g, Extended data Figure 5h,i). Thus, converging lines of evidence indicate that mFC neurons form mnemonic fields that track *task-progress* from specific behavioural steps.

#### 2- Distal prediction of behavioural choices

The SMB model posits that mFC neurons represent a memory of the animal’s past policy and provides a mechanistic explanation for this mnemonic function that we empirically validate above. The model also proposes a mechanism for how such neurons can be used to guide future behaviour. This is because the modules are proposed to be shaped by the structure of the task, in our case a loop of four rewards. This ring structure means modules can be used to repeat an effective policy once it is found (Figure 6a). If in trial N an animal makes a choice to visit a given behavioural step (e.g. early goal-progress location 2), then a bump of activity is initiated on the module that is anchored to this behavioural step at this point in the task (e.g. start of state B). This activity bump moves around the ring paced by progress in the task until it circles back close to the anchor point (e.g. towards the end of state A). This close-to-anchor activity biases the animal to return back to this same behavioural step in trial N+1 that it visited at the same point in the task in trial N (e.g. start of state B), thereby repeating the previous trial’s policy (Figure 6a). Thus, at any point in the sequence, the animal can choose among options for the next step in the maze by “listening” to the ring with the largest bump near its anchor point. The result is a sequential policy paced by the task periodicity. Crucially, the same memory buffers can be activated in a different sequence to encode a different task, without needing to build or bind new representations: thereby providing a programmable encoding of new task sequences.

**Figure 6 –.**
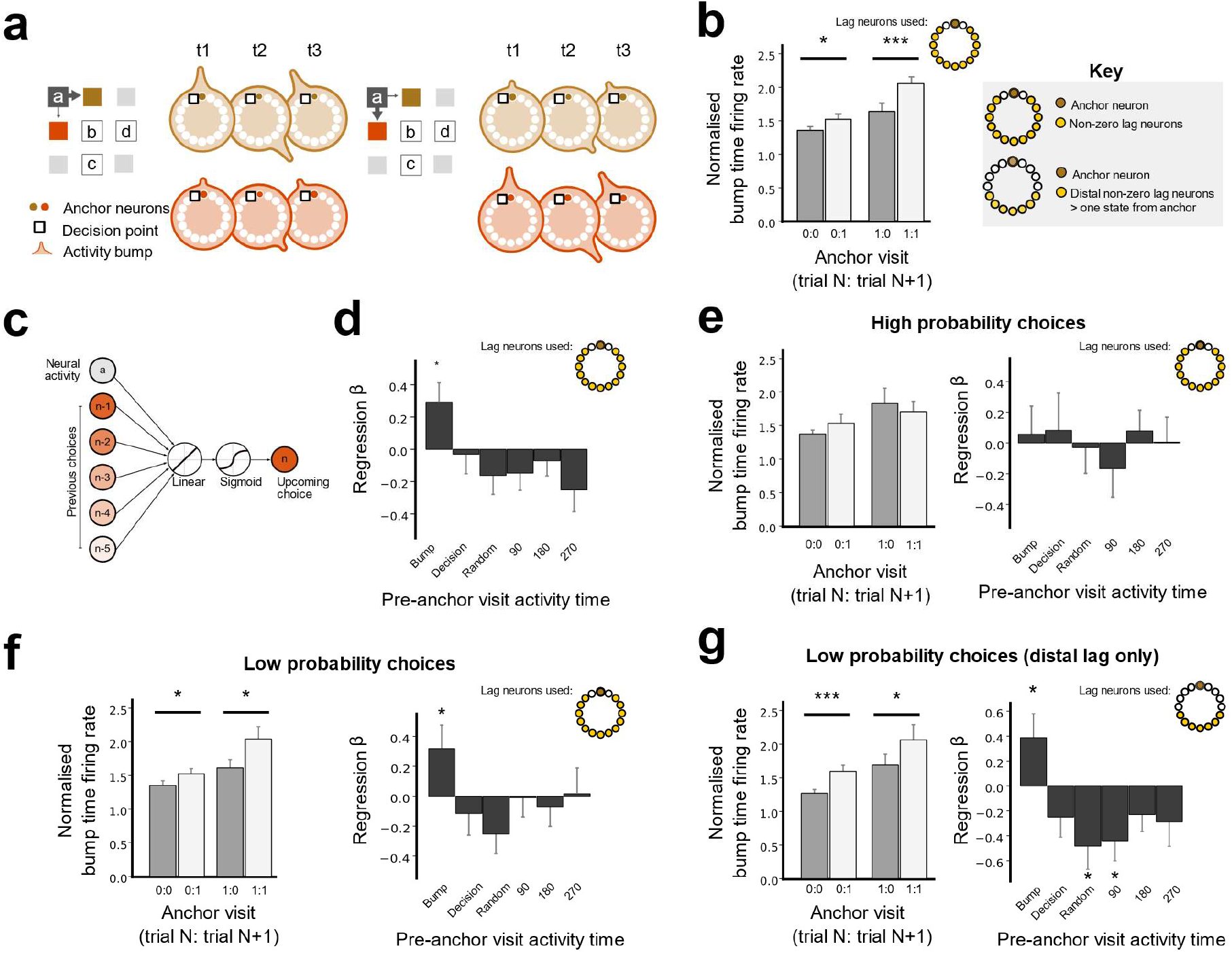
Medial Frontal activity predicts distal behavioural choices. a) Schematic showing distal prediction of animals choices from memory buffers. When the animal visits a goal-progress/place (t=1) in trial N, a bump of activity is initiated in the memory buffer that is anchored to this goal-progress/place. This bump travels around the buffer (e.g. t=2), paced by progress in the task. When the activity bump circles back to a point close to the anchor (t=3), it can be read out to bias the animal to return back to the same goal-progress/place in trial N+1 that was visited in the same task state in trial N. This read-out time defines a “decision point” that is specific for each memory buffer. Left: If, at t=3 in the example given, the bump on the buffer anchored to intermediate goal-progress in location 1 (brown square) is larger than that for the other option (intermediate goal-progress in location 4; red square) the animal will choose location 1. Right: Location 4 (red square) is chosen if the bump anchored to intermediate goal-progress in location 4 is larger at t=3. This choice could have been predicted from the bump sizes at an earlier time point (e.g. t=2) as the bump size will remain highly stable for the duration of a single trial, hence allowing distal prediction of choices from the memory buffers. b) Prediction of behaviour. Normalised firing rates of neurons during their “bump time”: i.e. the lag at which they are active relative to the anchor. X-axis labels denote visits to a goal-progress/place anchor in the current (N) and upcoming trial (N+1). For example a value of 0:1 means the anchor was not visited in trial N but visited in trial N+1. Bump time activity is higher before visits to the neuron’s anchor in trial N+1 whether the anchor was not visited in trial N (left) or when it was visited in trial N (right). Wilcoxon tests: Anchor not visited in trial N: n=78 sessions, statistic=1090, P=0.025. Anchor visited in trial N: n=72 sessions, statistic=666, P=2.76×10−4. In addition, an ANOVA on all data (N=72 sessions) showed a main effect of Past F=20.6 P=2.2×10−5, df1=1, df2=71, main effect of Future F=9.5 P=0.003, df1=1, df2=71 and no Past x Future interaction F=0.41 P=0.525, df1=1, df2=71. c) Design of logistic regression to assess effect of activity of the neuron on future visits to the anchor. To control for any autocorrelation in the choices of the a mouse, previous choices (binary values: 1s and 0s indicating visits) as far back as 5 trials in the past are added as regressors. Separate regressions are done for activity at different times: “bump time” (as in b); random times, decision time (the last 30 degrees of task space before the time where the animal could have visited the anchor of a given neuron, i.e. was one goal-progress-place away from it), and times shifted by 90 degree intervals relative to the neuron’s “bump time”. d) Regression coefficients are significantly positive for the bump time but not any of the other control times. T tests against 0: “bump time”: N=78 sessions, statistic=2.43, P=0.017, df=77; “decision time”: N=78, statistic=−0.26, P=0.792, df=77; “random time” N=78, statistic=−0.127, P=0.899, df=77; “90 degree shifted time” N=78, statistic=−1.34, P=0.183, df=77; “180 degree shifted time” N=78, statistic=−0.700, P=0.486, df=77; “270 degree shifted time” N=78, statistic=−1.86, P=0.066, df=77. e) Left: Distal prediction of behaviour to intermediate (non-rewarded) locations including choices to reward locations for only the <u>high probability</u> choices (choices that animals made with a probability in top half of maze transition probabilities during pre-task exploration). Bump time activity is indistinguishable before visits to the neuron’s anchor compared to non visits in trial N+1 whether the anchor was not visited in trial N (left) or when it was visited in trial N (right). Wilcoxon tests: Anchor not visited in trial N: n=59 sessions, statistic=728, P=0.236. Anchor visited in trial N: n=53, statistic=637, P=0.487. In addition, an ANOVA on all data (N=53 sessions) showed a trend towards a main effect of Past F=3.26 P=0.077, df1=1, df2=52, no main effect of Future F=0.026 P=0.871, df1=1, df2=52 and no Past x Future interaction F=1.60 P=0.211, df1=1, df2=52. Right: Regression coefficient is not significant for the bump time. T tests against 0: “bump time”: N=59 sessions, statistic=0.324, P=0.747, df=58; “decision time”: N=59, statistic=0.329, P=0.743, df=58; “random time” N=59, statistic=0.106, P=0.916, df=58; “90 degree shifted time” N=59, statistic=−0.918, P=0.362, df=58; “180 degree shifted time” N=59, statistic=0.611, P=0.544, df=58; “270 degree shifted time” N=59, statistic=0.052, P=0.959, df=58. f) Left: Distal prediction of behaviour to intermediate (non-rewarded) locations including choices to reward locations for only the l<u>ow probability</u> choices ((choices that animals made with a probability in bottom half of maze transition probabilities during pre-task exploration). Wilcoxon tests: Anchor not visited in trial N: n=64 sessions, statistic=744, P=0.048. Anchor visited in trial N: n=54, statistic=486, P=0.027. In addition, an ANOVA on all data (N=54 sessions) showed a main effect of Past F=5.66 P=0.021, df1=1, df2=52, a main effect of Future F=6.40 P=0.014, df1=1, df2=52 and no Past x Future interaction F=1.67 P=0.201, df1=1, df2=52. Right: Regression coefficients are significantly positive for the bump time but not any of the other control times. T tests against 0: “bump time”: N=64 sessions, statistic=2.03, P=0.047, df=63; “decision time”: N=64, statistic=−0.806, P=0.423, df=63; “random time” N=64, statistic=0.883, P=0.380, df=63; “90 degree shifted time” N=64, statistic=−0.079, P=0.937, df=63; “180 degree shifted time” N=64, statistic=−0.537, P=0.593, df=63; “270 degree shifted time” N=64, statistic=0.100, P=0.920, df=63. g) Prediction of behaviour using neurons anchored at distal points (one state away or more) from the anchor for only the <u>low probability</u> choices (choices that animals made with a probability in bottom half of maze transition probabilities during pre-task exploration). Normalised firing rates of neurons during their “bump time”: i.e. the lag at which they are active relative to the anchor. Bump time activity is higher before visits to the neuron’s anchor in trial N+1 whether the anchor was not visited in trial N (left) or when it was visited in trial N (right). Wilcoxon tests: Anchor not visited in trial N: n=67, statistic=463, P=2.41×10^−5^. Anchor visited in trial N: n=51, statistic=430, P=0.029. In addition, an ANOVA on all data (N=51 sessions) showed a main effect of Past F=5.29 P=0.026, df1=1, df2=50, no main effect of Future F=10.19 P=0.002, df1=1, df2=50 and no Past x Future interaction F=7.73×10^−4^ P=0.978, df1=1, df2=50. Regression coefficients are significantly positive for the bump time but not any of the other control times. T tests against 0: “bump time”: N=67 sessions, statistic=2.02, P=0.048, df=66; “decision time”: N=67, statistic=−1.54, P=0.128, df=66; “random time” N=67, statistic=−2.60, P=0.011, df=66; “90 degree shifted time” N=67, statistic=−2.73, P=0.008, df=66; “180 degree shifted time” N=67, statistic=−1.65, P=0.103, df=66; “270 degree shifted time” N=67, statistic=−1.46, P=0.148, df=66. All error bars represent the standard error of the mean

This model makes a unique prediction: the choice an animal will make to visit a particular maze location should be predictable from the activity of neurons anchored to the possible choices several steps and 10s of seconds before the animal actually makes a decision. This is because the bump of activity is constrained to move around a given ring. Hence, bump size at an early point in the trial should strongly correlate with its size just before the anchor point (Figure 6a). The time point just before the anchor point is the “decision time” for a given ring where the output neurons of the ring can be read out to bias the animal’s choice towards the behavioural step represented by the anchor (Figure 6a). The time at which the putative bump of activity passes through a given neuron (the neuron’s “bump-time”) reflects the neuron’s task-space lag from the behavioural step to which its anchored: i.e. its position on the ring relative to the anchor (Figure 4a, Figure 6a). The SMB model proposes that activity restricted to this “bump-time” can be used to predict whether an animal will return back to this same behavioural step at the same point in the next trial. Because a trial is on average ~30 seconds in length, this means an animal’s choices can be predicted 10s of seconds before they are made. To test this prediction, we related the trial-by-trial “bump-time” activity of neurons at non-zero lag to their anchor with subsequent visits to this same anchor. Intriguingly, while controlling for previous choices, we found that the “bump-time” activity of neurons was higher before animals visited the neurons’ anchor (Figure 6b). To investigate this further, we ran a logistic regression on the trial by trial activity of neurons at non-zero lag from their anchor to predict upcoming behavioural choices by the animal. To control for the autocorrelation in animals’ behaviour, we regressed out previous choices going all the way back to 5 trials in the past (Figure 6c). We found that mFC neurons significantly predicted future choices only when taking their activity at the correct “bump-time”. Crucially, activity of a given neuron at the “wrong” times, whether at random timepoints, times shifted by 90 degree intervals relative to the “bump-time” to preserve goal-progress tuning, or even at the “decision time” itself, did not predict subsequent choices (Figure 6d). This prediction held even when only considering choices to intermediate, non-rewarded locations (Extended data figure 6a).

Notably, animals are not performing optimally (Figure 1d). Whilst performance is well above chance, they continue to be biased by pre-existing policies that existed before they learned any ABCD task (Extended data Figure 1c). We reasoned that if the frontal cortex exerts goal directed control^21^, then we might only be able to predict choices where animals take actions that go against their pre-existing biases. We therefore divided choices into high and low probability ones, based on the probability animals made a given choice in an exploration session on the same maze but before the animals had been exposed to any ABCD task. Intriguingly, we found that frontal activity was not predictive of high probability choices (Figure 6e). In contrast, mFC activity predicted low probability choices (Figure 6f). This is in line with the view that frontal neurons are preferentially implicated in altering behaviour against stereotyped policies^21^. Moreover, “bump-time” activity of neurons with mnemonic fields that are lagged by more than one state away from the anchor also predicted choices (Extended data figure 6b). This again was seen for low (Figure 6g) but not high probability choices (Extended data figure 6c). Overall these results, showing a distal prediction of behavioural choices, empirically validate the SMB model as a mechanism for the retrieval of task-paced policies by the mFC.

#### 3- Internally organised memory buffers

Our findings so far support the view that mFC modules invariantly track task-progress from different behavioural steps, and use this memory to retrieve a sequential policy that is paced by the task’s periodicity. To retrieve such task-paced sequences, the SMB model proposes that memory buffers are shaped by the structure of the task. In the ABCD task, the structure is a loop consisting of 4 rewarded goals (4 goal-progress cycles). Thus the neuronal state space should curve back onto itself after 4 goals, creating an internally organised ring structure. While the policy retrieval results (Figure 6; Extended data figure 6) are consistent with this view, we sought a more direct test of this task-structuring hypothesis. To this end, we investigated whether pairwise coactivity during pre-task sleep reflects a ring-like neuronal state space. We regressed the circular distance between pairs of neurons that share the same anchor, and hence belong to the same module, against their cross-correlation during sleep (Figure 7a). This was done while co-regressing the forward distance from the anchor: the distance between two neurons in a state space shaped into a line that begins with the anchor (Figure 7a). We found that regression coefficient values were significantly negative for circular distance, while controlling for forward distance, indicating that neurons closer on a circular state space (smaller neuron-neuron distances on a ring) are more coactive (Figure 7b). These results held while controlling for pairwise spatial map similarity and goal-progress-tuning proximity between neurons, which were added as co-regressors. Forward distance did not predict co-activity when controlling for circular distance (Figure 7b). Moreover, we isolated pairs of neurons that had systematically opposed circular and forward distances (i.e. pairs of neurons separated by a forward distance of 180 degrees or more). Regressing circular distance against sleep coactivity for only those neuron pairs also showed a significantly negative regression coefficient, further supporting a ring-shaped state space (Figure 7c). In addition, pairs of consistently anchored neurons that share the same anchor showed significantly more negative regression coefficient values than neuron pairs across anchors (Figure 7d). This was true both for pre-task and post-task sleep (Figure 7d). Taken together, these findings provide offline evidence that mFC modules are internally organised by the structure of the task.

**Figure 7 –.**
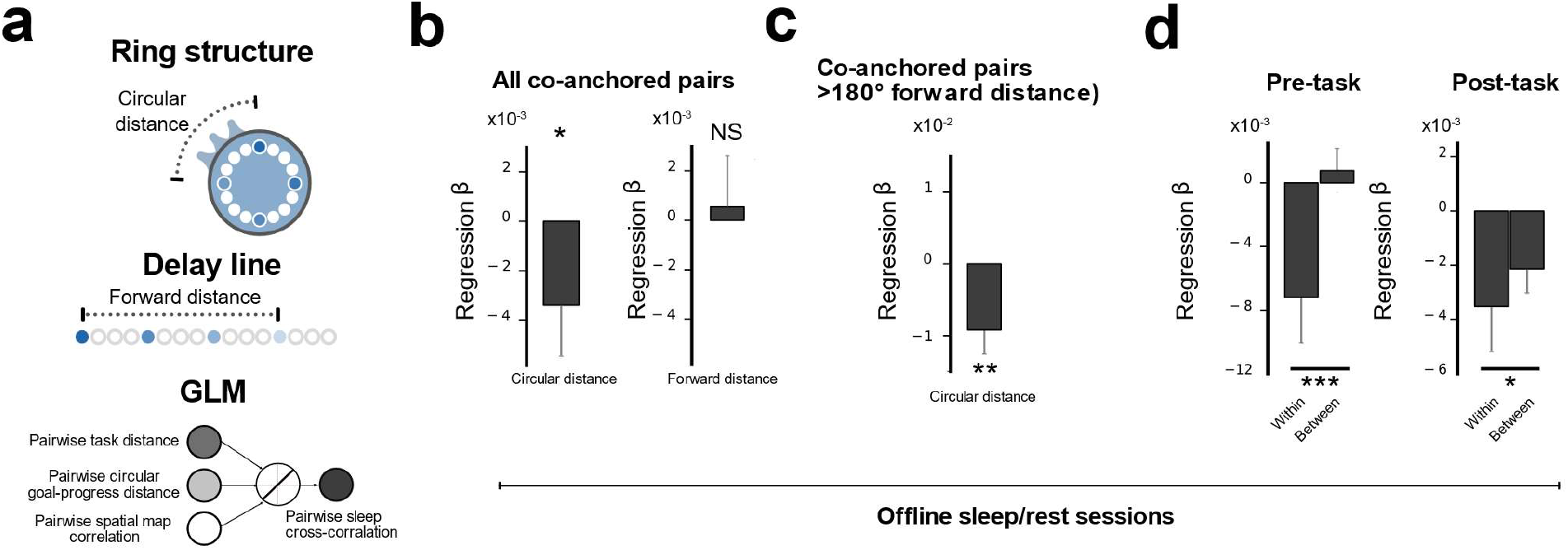
Offline activity of mFC neurons is internally organised by the Task structure. a) Top: Schematic showing potential neuronal state spaces - if neurons are arranged on a ring, then circular distance is a better description of how close two neurons are in state space than forward distance relative to the anchor. Conversely, if neurons lie on a delay line, forward distance is a better description of neuron-neuron co-firing relationships. Bottom: Schematic showing the inputs and outputs of linear regression model relating pairwise task-distance (either circular or forward distance) with coactivity during sleep while regressing out pairwise goal-progress tuning distance and spatial map similarity. b) Regression coefficient values for circular (left) or forward (right) distance regressed against sleep cross-correlation for co-anchored neurons - T test relative to 0 (One-tailed): circular distance: N=612 pairs t=−1.656 P=0.049; forward distance: t=0.266 P=0.395, df=612 c) Regression coefficient values for circular distance for only co-anchored pairs of neurons separated by a forward distance of 180 degrees or more in task space: T test relative to 0 (One-tailed): N=149 pairs t=−2.770 P=0.003, df=148 Note that forward distance is the exact inverse of circular distance for forward distances of 180 degrees or higher, and hence forward distance regression coefficients will be exactly the same magnitude, but opposite sign, to circular distance regression coefficients. d) Regression coefficient values for circular distance against sleep cross-correlation using pairs of neurons consistently anchored to the same anchor (within) vs pairs with different anchors (between). Regression coefficient values were significantly more negative for within compared to between comparison for pre-task (left) and post-task (right) sleep. Two-tailed unpaired T test: Pre-task: t = 7.7216, N=268 pairs (within), 2722 pairs (between), df = 2988, P<0.0001; Post-task: N=256 pairs (within), 2774 pairs (between), df = 3028, t = 2.3145,P=0.0207 All error bars represent the standard error of the mean

## Discussion

Our findings identify a cellular algorithm for mapping abstract behavioural structure. We found that mice can learn an abstract structure organising multiple goals and use it to rapidly learn complex behavioural sequences. The large majority of mFC neurons tiled progress-to-goal, generalising across behavioural sequences of different distances, times and locations. This goal-progress tuning was further elaborated to form representations shaped by the overall structure of the task, a four-goal loop in our case. The use of goal-progress tuning as a scaffold for building task-structure representations ensured that mFC neuronal dynamics evolved as a function of true progress in the overall task as defined by the goals, rather than other physical dimensions. Using goal-progress-tuned neurons also meant that frontal representations tracked task progress not only from each goal location but also from intermediate behavioural steps *en route* between goals. This allowed encoding long sequences of behaviours that were parsed by the task structure, guided primarily by the network dynamics of pre-learned memory buffers. The resulting algorithm provides a unified account of mFC roles in schema formation and sequence memory, by internally organising mnemonic activity according to an abstract structure shared by many tasks. It further suggests a general mechanism for building behavioural schema by using “goal-progress” neurons as a building block that can be sculpted to represent the structure of any complex task.

The SMB algorithm we discover here can potentially reconcile a number of findings concerning the mFC. Lesions to the Frontal cortex across species affect the sequencing of complex, goal-directed behaviours^37^. Here we show that if mnemonic representations resembling those implicated in sequential working memory^22^ are shaped by an abstract task structure, they could explain another key function of the mFC: the formation of generalised task schema^12^. Moreover, the differences in the representational logic employed at each level of the task hierarchy seen in our study could potentially reconcile seemingly disparate views on task structure generalisation. At the level of tracking progress towards a single goal, we find that neurons tile this space using fixed sequences that generalise across different behavioural trajectories. These goal-progress cells are similar in spirit to neurons in the mFC that encode general task states regardless of sensorimotor specifics^11,28–31^. However, at the level of tracking task-progress in a sequence comprising multiple goals, network dynamics are not universally coherent across all neurons, but are instead apportioned into modules, each tracking a memory of a particular behavioural step. Such representational logic is consistent with recent empirical findings in sequence working memory tasks^22,38^ and tasks where tracking previous choices and/or reward history is necessary for optimal behaviour^23,24^ as well as computational work on recurrent neural networks trained to generalise across tasks^35^. Moreover, while such SWM tasks are not explicitly circular, maintenance of the experienced sequence in the delay period might be implemented via similar ring attractor dynamics. These dynamics would later guide retrieval of the sequence in the same way we describe here, through activating anchor neurons representing the next step in the sequence via simulation rather than direct experience. More generally, we propose that invariant representations may track progress towards a single goal while stimulus-specific working-memory-like representations are used for tracking progress in a complex task comprising multiple goals of equal valence.

How does the organisation of neurons in the mFC mapping “behavioural structure” compare with the cellular bases of cognitive maps of “world structure”? Like the grid-cells that underlie world-structure mapping, frontal neurons are internally organised according to the structure they map. Neurons maintain this structure when the contents of the sequence they are mapping change in new tasks in mFC (Figure 3) and in new spatial contexts in mEC^36^, and even in the absence of any structured sensory input during sleep (Figure 7 for mFC; ^39,40^ for mEC). We also note the modularity that underlies both types of maps (Figure 3 for mFC; ^36^ for mEC), which may be a general feature of cortical representations that allows mapping spaces from different reference points^41^. There are also key differences. One difference relates to the velocity signal driving dynamics in both maps. By virtue of needing to map behavioural space rather than physical space, activity bumps along attractors representing behavioural structure evolve as a function of progress towards a goal (Figure 2), while grid cells use the velocity of the animal in physical space to track its spatial position^42^. This basic goal-progress scaffold in the Frontal cortex enables building a representation that evolves as a function of task stages, allowing animals to reliably map true task-progress independently of other variables such as elapsed time or distance. Another apparent difference relates to how maps are used to guide flexible behaviour in new contexts. For maps of world structure, convergent input from medial Entorhinal grid cells and sensory input from the lateral Entorhinal cortex into the Hippocampus is thought to give rise to the canonical “place cell” in a new spatial context^43^. This grounding of abstract, relational “world-structure” knowledge to concrete experiences allows us to navigate, path integrate and infer new routes to goals^3,4,44,45^. Conversely, by virtue of having fixed anchoring to concrete behavioural steps, frontal neurons store new behavioural sequences in the dynamics of the network, without needing access to an external memory and new plasticity. This offers a programmable solution to mapping new sequences, which avoids the need for associative binding on new tasks. Intriguingly, a recent study suggests that similar fixed anchoring of grid-cells to landmarks may allow rapid, plasticity-free mapping of new spatial environments^46^, akin to what we report here for frontal maps of behavioural structure. How associative and network dynamics-based solutions compare computationally, how they coexist in the same and distinct circuits and when they are used remain open questions. We address some of these computational questions in a recent theoretical study^47^.

The programmable solution employed to map behavioural structure in expert animals places the burden of plasticity on sculpting attractor dynamics when learning early tasks. How is such learning achieved? A potential answer comes from considering that memory buffers representing the abstract task structure are expressed on top of lower level goal-progress tuning. Goal-progress-tuned neurons may have been initially part of task-naive goal-progress sequences that then became sculpted, through learning of many examples, to form structured memory buffers. These naive goal-progress sequences may develop first to map sequences to any behavioural goal independently of any higher order structure organising the goals. Input from the Hippocampus and/or medial Entorhinal cortex could provide spatial information to the mFC^48^ to generate the conjunctive goal-progress/place anchors. Plasticity at recurrent mFC-thalamus-Basal ganglia loops could then allow learning task-structured memory buffers that track task-progress from these anchors^49,50^. Goal-progress sequences could therefore provide an inductive bias which facilitates the formation of a schema encoding any task-structure in the mFC. Monitoring and manipulating mFC neurons and their inputs during learning of the ABCD task and equivalent hierarchical learning paradigms will allow understanding the inductive biases, learning rules and circuit mechanisms that generate maps of behavioural structure.

## Extended Data Figures

**Extended Data Figure 1.**
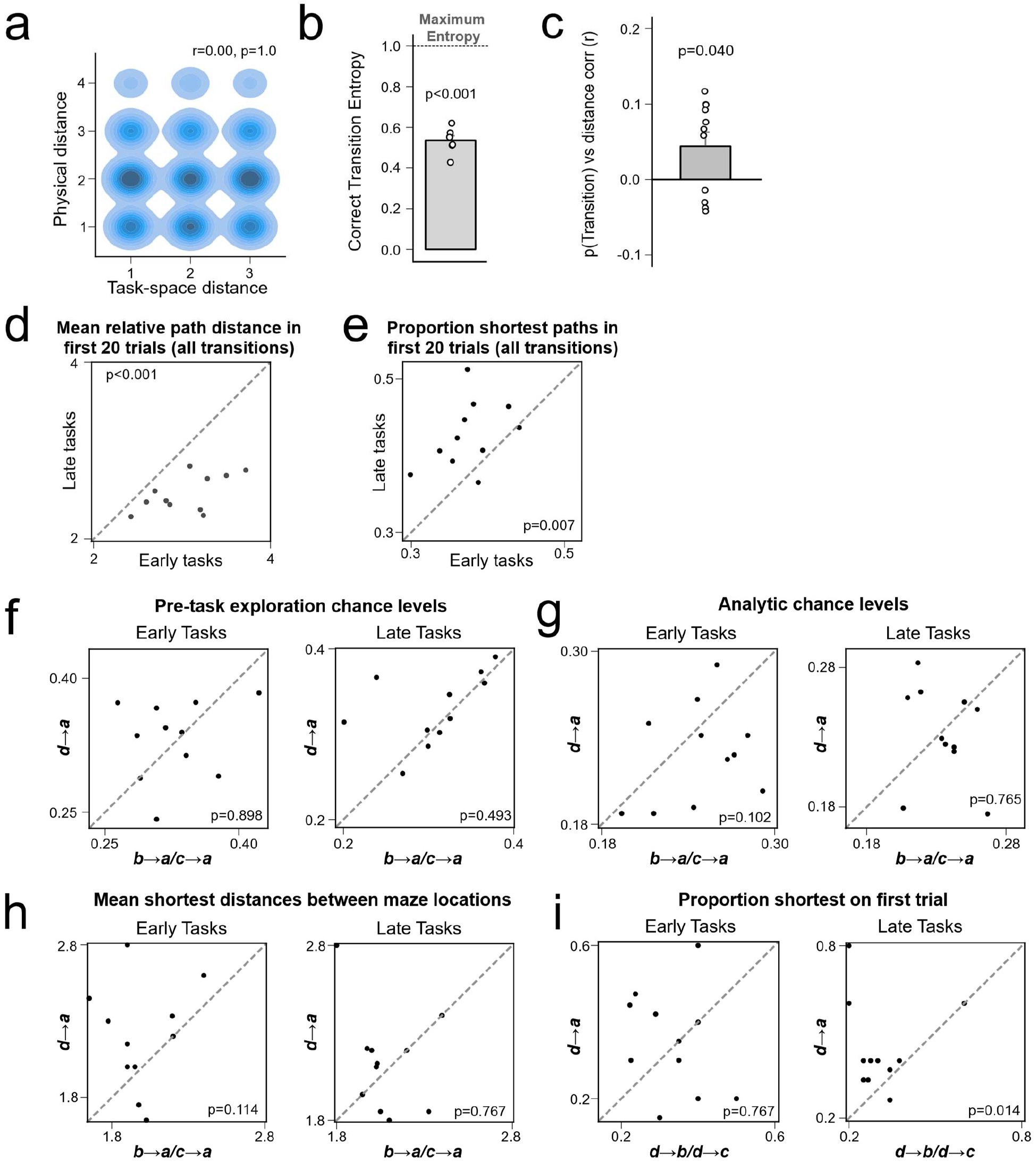
a) Tasks were designed such that task space and physical space are orthogonal to each other. A kernel density estimate plot showing that distances between reward locations in task space (how many states are between the rewards) are not correlated with the physical distances in the maze. Data points are individual tasks. Pearson correlation: r=0.00 P=1.0 b) Animals used stereotyped routes when taking the shortest route to a goal. The entropy of correct transitions taken is lower than expected if animals took all shortest routes equally. T-test against 1 – N=11 animals, statistic=−20.38, P=9.07×10^−7^, df=10 c) Suboptimal performance was associated with persisting behavioural biases from before exposure to the task. Y-axis shows the r value calculated from a correlation between the mean relative path distance taken between goals and the probability the transitions within this trajectory would have been taken when the animal was naive to any ABCD task (when the animal explored the arena before any rewards or tasks were presented). A net positive correlation indicates that when animals take longer routes (i.e. perform less optimally) they take these routes through transitions that they were more likely to take before exposure to any ABCD task. T-test against 0 – N=11 animals, statistic=2.36, P=0.040, df=10 d) Mean relative path distance travelled by the mice between goals in the first 20 trials of early vs late tasks. Wilcoxon test N=11 animals, Statistic=0.0, P=9.77×10−4 e) Mean proportion of transitions where one of the shortest routes was taken in the first 20 trials of early vs late tasks. Wilcoxon test N=11 animals, Statistic=4.0 P=0.007 f) No difference in the empirical chance levels (baseline transition probabilities calculated when animals explored the maze before experiencing any ABCD tasks: see Methods under “Behavioural Scoring”) between ***d*** to ***a*** and ***c***-to-***a***/***b***-to-***a*** transitions on the *first* trial in early (left) and late (right) tasks. Wilcoxon test; Early tasks: N=11 animals, statistic=31.0, P=0.898; Late tasks: N=11 animals, statistic=23.0, P=0.413 g) No difference in the analytical chance levels (see Methods under “Behavioural Scoring”) between ***d*** to ***a*** and ***c***-to-***a***/***b***-to-***a*** transitions on the *first* trial in early (left) and late (right) tasks. Wilcoxon test; Early tasks: N=11 animals, statistic=14.0, P=0.102; Late tasks: N=11 animals, statistic=29.0, P=0.765 h) No difference in the shortest physical maze distances between ***d*** to ***a*** and ***c***-to-***a***/***b***-to-***a*** transitions on the *first* trial in early (left) and late (right) tasks. Wilcoxon test; Early tasks: N=11 animals, statistic=12.0, P=0.114; Late Tasks: N=11 animals, statistic=20.0, P=0.767 i) Zero-shot inference on the first trial of late tasks is associated with animals returning from ***d*** to ***a*** more often than ***d***-to-***b*** or ***d***-to-***c***. The proportion of tasks in which animals took the most direct path from ***d***-to-***a*** on the *first* trial is compared to the same measure but for premature returns from ***d****-*to-***b*** and ***d****-*to-***c*.** Early tasks are shown on the left and late tasks on the right. Wilcoxon test; Early tasks: N=11 animals, statistic=20.0, P=0.767; N=11 animals, Late tasks: statistic=6.0, P=0.014 All error bars represent the standard error of the mean

**Extended Data Figure 2.**
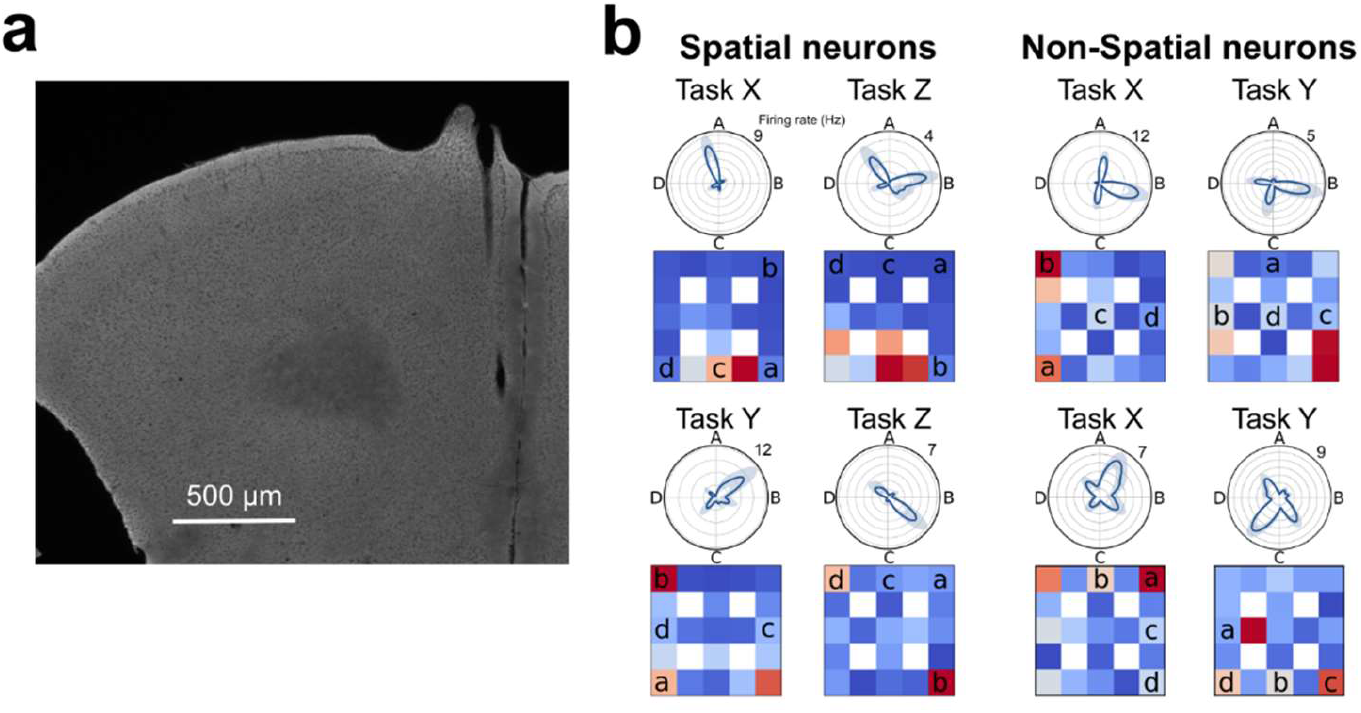
a) Coronal slice from an implanted mouse showing silicon probe track terminating in the prelimbic region of mFC. b) Polar plots of task tuning and spatial maps for four example neurons that are tuned to both goal-progress and state. Each neuron is plotted across two tasks to illustrate spatial tuning (left two neurons) and lack thereof (right two neurons).

**Extended Data Figure 3.**
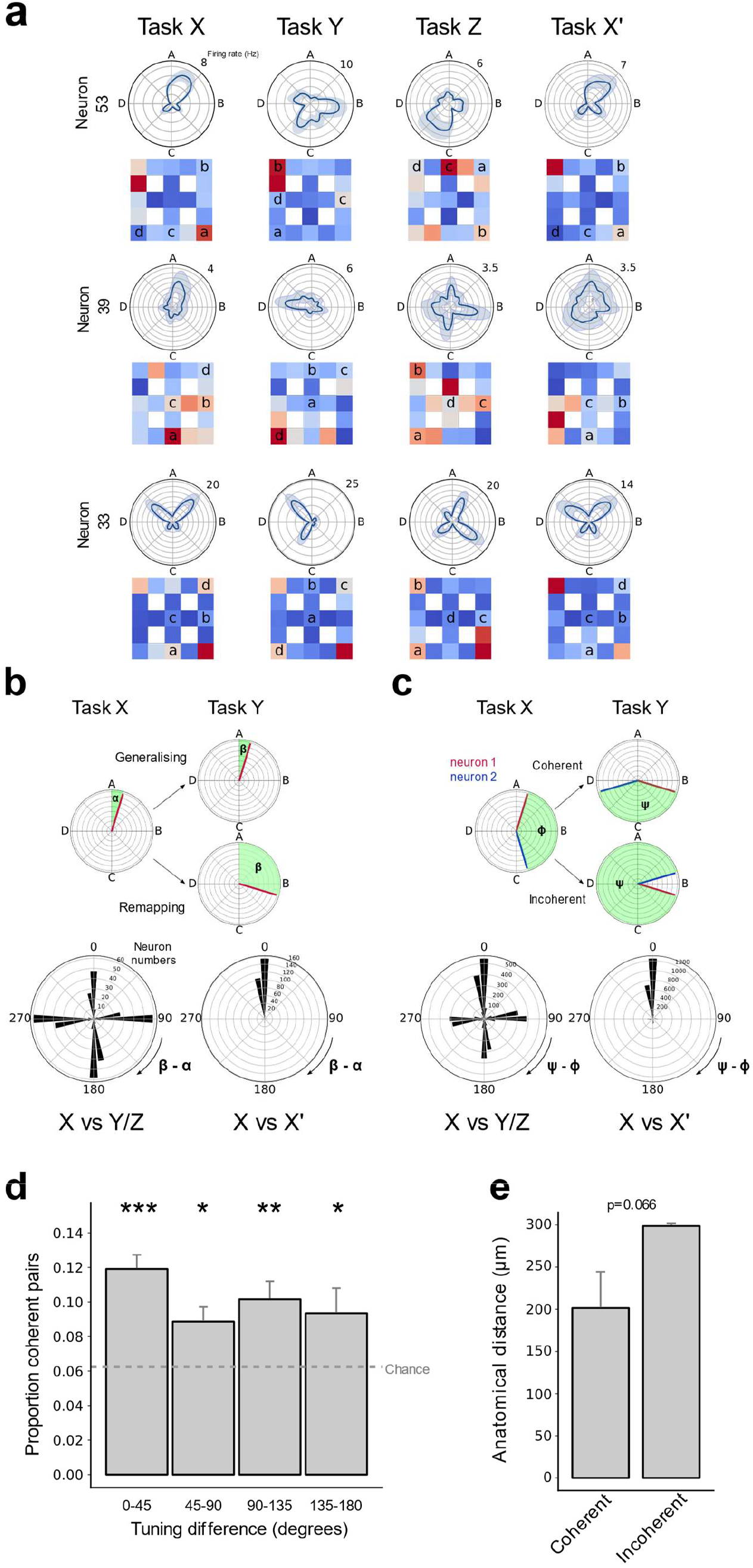
a) Three example state-tuned neurons remapping across tasks. The top two neurons remap in a way that is not related to their spatial maps in any given task. The bottom neuron remaps in accordance to its spatial map. b) Remapping of non-spatial neurons. Top: Quantifying the difference in tuning angles for the same neuron across tasks (top). Bottom left: Polar histograms show that non-spatial state-tuned neurons remap by angles close to multiples 90 degrees, as a result of conserved goal-progress tuning and the 4 reward structure of the task. No clear peak at zero is seen relative to the other cardinal directions when comparing sessions spanning separate tasks (Two proportions test against a chance level of 25% N=432 neurons; mean proportion of generalising neurons across one comparison (mean of X vs Y and X vs Z)=21%, z=7.01, P=2.33×10−12: i.e. significantly lower than chance). Bottom right: Non-spatial state-tuned neurons maintain their state preference across different sessions of the same task (bottom right). Two proportions test against a chance level of 25% N=324 neurons; proportion generalising=84%, z=20.3, P<0.001). c) Coherence of non-spatial neuron pairs. Top: Quantifying difference in relative angles between neurons across tasks. Bottom: Only non-spatial state-tuned neurons are shown. Bottom left: Polar histograms show that the proportion of coherent pairs of non-spatial state-tuned neurons (comprising the peak at zero) is higher than chance but far from 1, indicating that the whole population does not rotate coherently (Two proportions test against a chance level of 25% N=3946 pairs; mean proportion of coherent neurons across one comparison (mean of X vs Y and X vs Z)=31%, z=29.8, P<0.001)). Bottom right: As expected from panel b, the large majority of non-spatial state-tuned neurons keep their relative angles across sessions of the same task (X vs X’; Two proportions test against a chance level of 25% N=2805 pairs; proportion coherent=75%, z=53.6, P<0.001). d) Proportion of coherent pairs per recording day (pairs of state-tuned neurons where the relative angle doesn’t change by more than 45 degrees across *both* X to Y and X to Z comparisons) relative to all pairs across different pairwise task space angles. T tests with Holm-Sidak correction against chance level of 1/16 (probability of neuron pair rotating coherently across two comparisons (i.e. 0.25^2^)): N=21 recording days, pairwise circular distance difference: 0-45 degrees statistic=6.89, P=4.33×10^−6^, df=20; 45-90 degrees statistic=3.10, P=0.011, df=20; 90-135 degrees statistic=3.74, P=0.004, df=20; 135-180 degrees statistic=2.14, P=0.044, df=20. e) Coherent pairs show a trend towards being closer anatomically than incoherent pairs. Independent T test: N=4138 pairs, statistic=−1.84, P=0.065, df=4136 All error bars represent the standard error of the mean

**Extended Data Figure 4.**
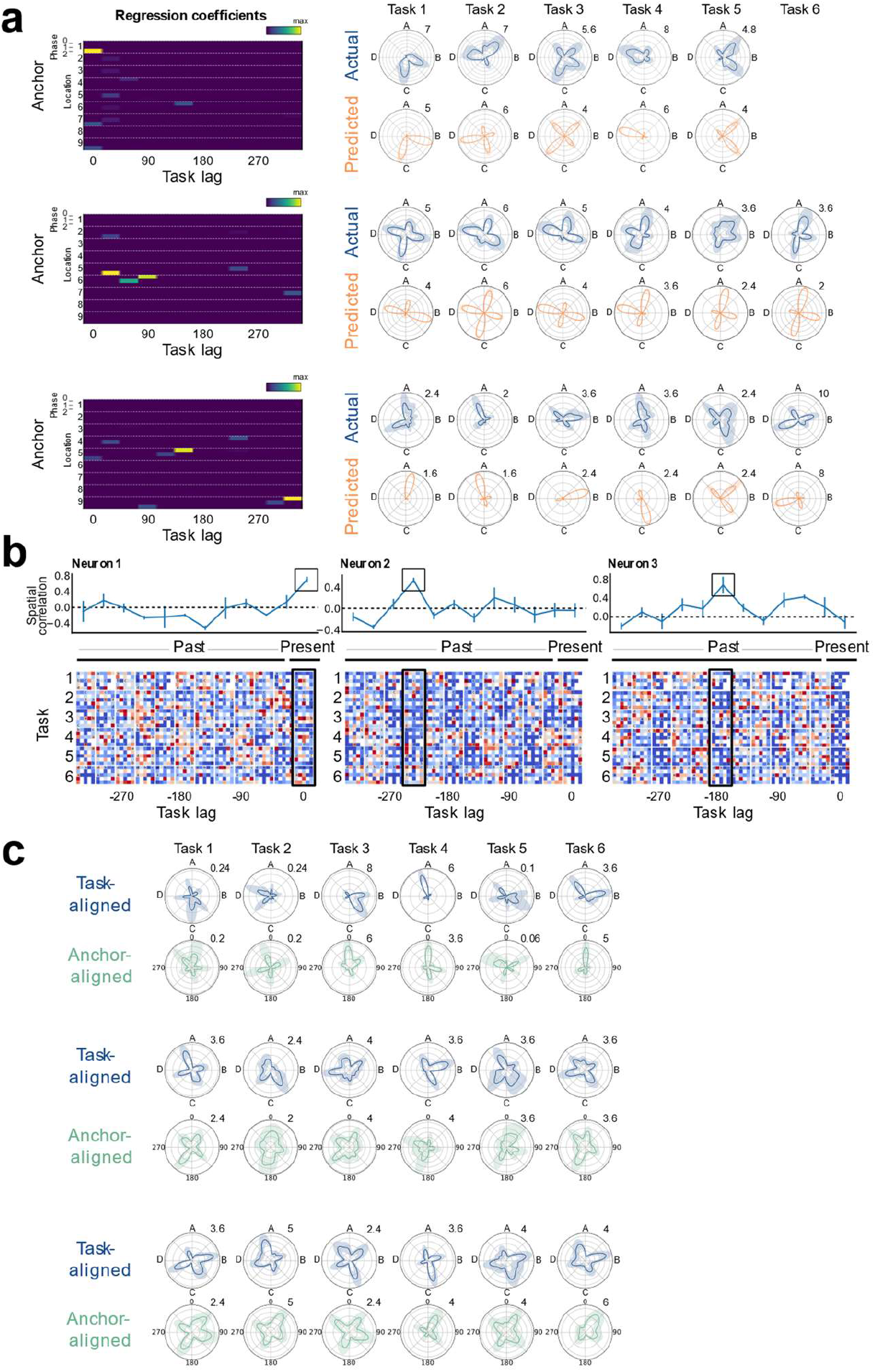
a) Regression analysis reveals neurons with mnemonic fields lagged in task space from a given goal-progress/place anchor (bottom two neurons), alongside neurons directly tuned to a goal-progress/place (top neuron). The regression coefficients are shown on the left with the actual (blue) and predicted (orange) activity of the neurons shown on the right. b) Lagged spatial field analysis. Example plots showing spatial maps for 3 neurons. Each row represents a different task and each column a different lag in task space. Bottom: Activity of each neuron is plotted as a function of the animal’s current location (far right column for each cell) and at successive task space lags in the past for the remaining columns. Top: the correlation of spatial maps across tasks at each lag. The neuron on the left is an anchor cell (goal-progress/place cell), as seen by the correlation peak at zero lag, while the middle and right-most neurons are neurons lagged by 240 and 180 degrees from their anchors respectively. c) Single anchor alignment analysis. Top (blue) plots for each neuron shows activity aligned by the abstract states (with the dashed vertical line at zero representing reward *a*, i.e. the start of state A; going clockwise, the remaining dashed lines represent reward locations ***b***, ***c*** and ***d***, and hence the starts of states B C and D respectively). Neurons appear to remap in task space across tasks. Bottom (green) plots for each cell show that it is possible to find a goal-progress/place anchor that consistently aligns neurons across tasks (the zero line corresponds to visits to the goal-progress/place anchor). Top neuron has a peak at zero relative to the anchor, while the middle and the bottom neurons peak at 200 degrees and 45 degrees from their anchors respectively.

**Extended Data Figure 5.**
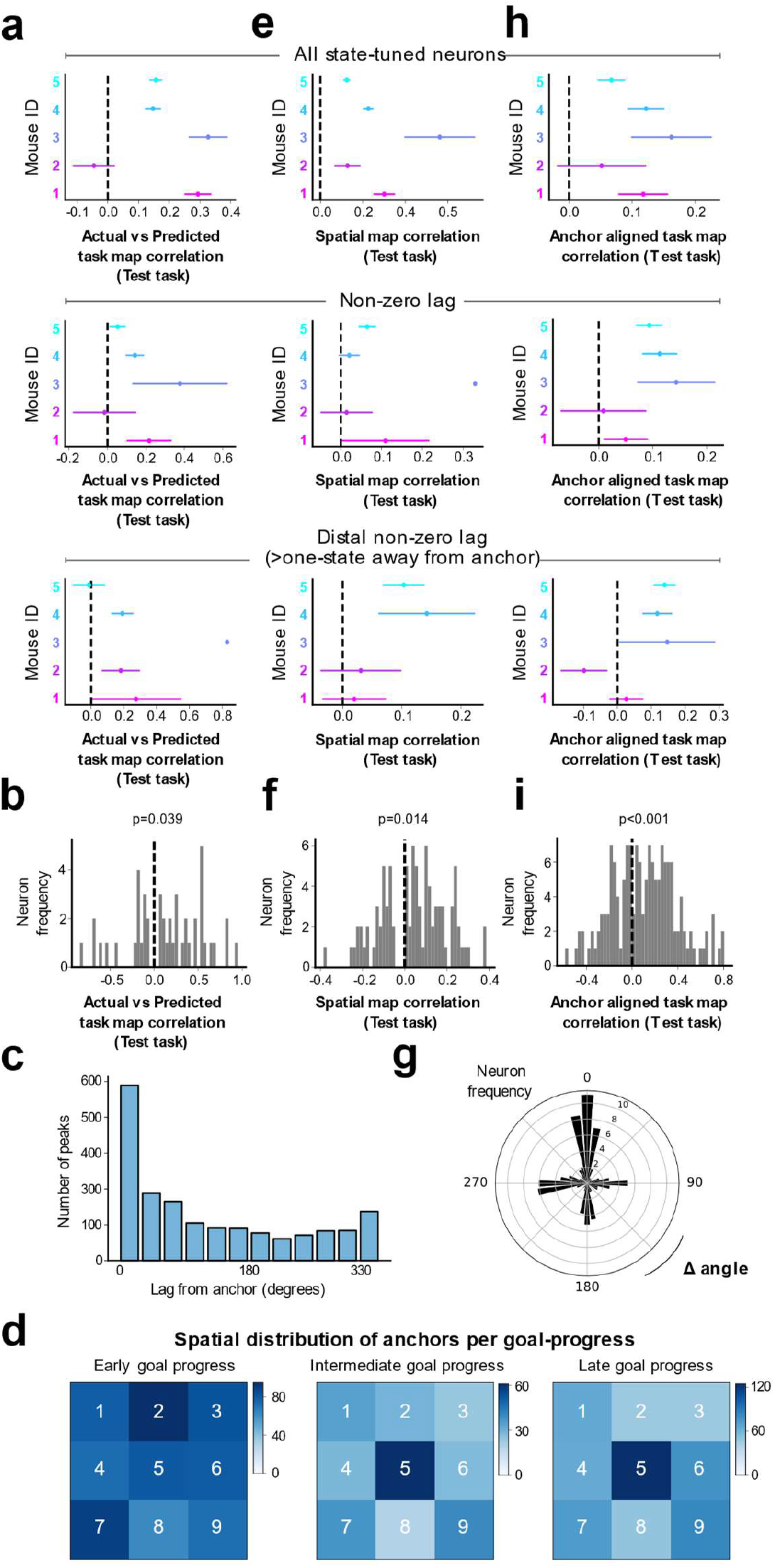
a) Mean, per mouse distribution of cross-validated correlation values between model-predicted (from training tasks) and actual activity (from a left out <u>test task</u>) for: Top: All state-tuned neurons, Middle: non-zero lag state-tuned neurons (30 degrees or more away from anchor), Bottom: distal non-zero lag state-tuned neurons (90 degrees or more away from anchor). One-tailed binomial test against chance (chance being mean values equally likely to be above or below 0): All state-tuned neurons: 4/5 mice with mean positive correlation P=0.1875; Non-zero lag state-tuned neurons: 4/5 mice with mean positive correlation P=0.1875; Distal Non-zero-lag state-tuned neurons: 4/5 mice with mean positive correlation P=0.1875). b) Histogram showing the right shifted distribution of mean cross-validated correlation values between model-predicted (from training tasks) and actual activity (from a left out <u>test task</u>) for only non-zero lag state-tuned neurons with one of the top 3 maximum regression coefficient values a whole state (90 degrees) or more either side of the anchor. To avoid contamination due to potential residual spatial-tuning, only regression coefficient values more than 90 degrees in task space either side of the anchor point are used for the prediction. T test against 0: N=46 neurons, statistic=2.13, P=0.039, df=45. c) Distribution of task space lags from anchor for all state-tuned neurons. d) 2D histograms showing spatial and goal-progress distributions of anchors. The colour bar represents the number of neurons anchored to each maze location at each goal-progress (maze repeated 3 times to display results for early, intermediate and late goal-progress anchors). e) Mean, per mouse distribution of cross-validated spatial correlations between spatial maps at the preferred lag (from training tasks) and the spatial map at this lag from a left out <u>test task</u> for: Top: All state-tuned neurons Middle: non-zero-lag state-tuned neurons (30 degrees or more away from anchor) Bottom distal non-zero-lag state-tuned neurons (90 degrees or more away from anchor). One-tailed binomial test against chance (chance being mean values equally likely to be above or below 0): All state-tuned neurons: 5/5 mice with mean positive correlation P=0.03125; Non-zero lag state-tuned neurons: 5/5 mice with mean positive correlation P=0.03125; Distal Non-zero lag state-tuned neurons: 4/4 mice with mean positive correlation P=0.0625). f) Histogram showing the right shifted distribution of the mean cross-validated spatial correlations between spatial maps at the preferred lag in training tasks and the spatial map at this lag from a left out <u>test task</u> for only non-zero-lag state-tuned neurons with spatial correlation peaks a whole state (90 degrees) or further either side of zero-lag. T test against 0: N=90 neurons, statistic=2.49, P=0.015, df=89. g) Polar histogram showing the cross-validated alignment of non-zero-lag neurons by their preferred goal-progress/place (calculated from training tasks) in a left-out test task. Only neurons with a lag of 90 degrees (one state) or more either side of their anchor are shown. Two proportions test against chance (25%): Proportion generalising=39.2%; N=154 neurons, z=7.23, P=4.98×10^−13^ h) Mean, per mouse distribution of cross-validated task map correlations between neurons aligned to their preferred goal-progress/place anchor (from training tasks) and the task map aligned to this goal-progress/place from a left out <u>test task</u> for: Top: all state-tuned neurons Middle: non-zero-lag state-tuned neurons (30 degrees or more away from anchor) Bottom distal non-zero-lag state-tuned neurons (90 degrees or more away from anchor). One-tailed binomial test against chance (chance being mean values equally likely to be above or below 0): All neurons: 5/5 mice with mean positive correlation P=0.03125; Non-zero-lag neurons: 5/5 mice with mean positive correlation P=0.03125; Distal Non-zero-lag neurons: 4/5 mice with mean positive correlation P=0.188). i) Histogram showing the right shifted distribution of the mean cross-validated task map correlations between neurons aligned to their preferred goal-progress/place anchor (from training tasks) and the task map aligned to this goal-progress/place from a left out <u>test task</u> for only non-zero-lag state-tuned neurons with a lag of 90 degrees or more either side of their anchor. T test against 0: N=154 neurons, statistic=4.59, P=9.38×10^−6^, df=153. All error bars represent the standard error of the mean

**Extended Data figure 6.**
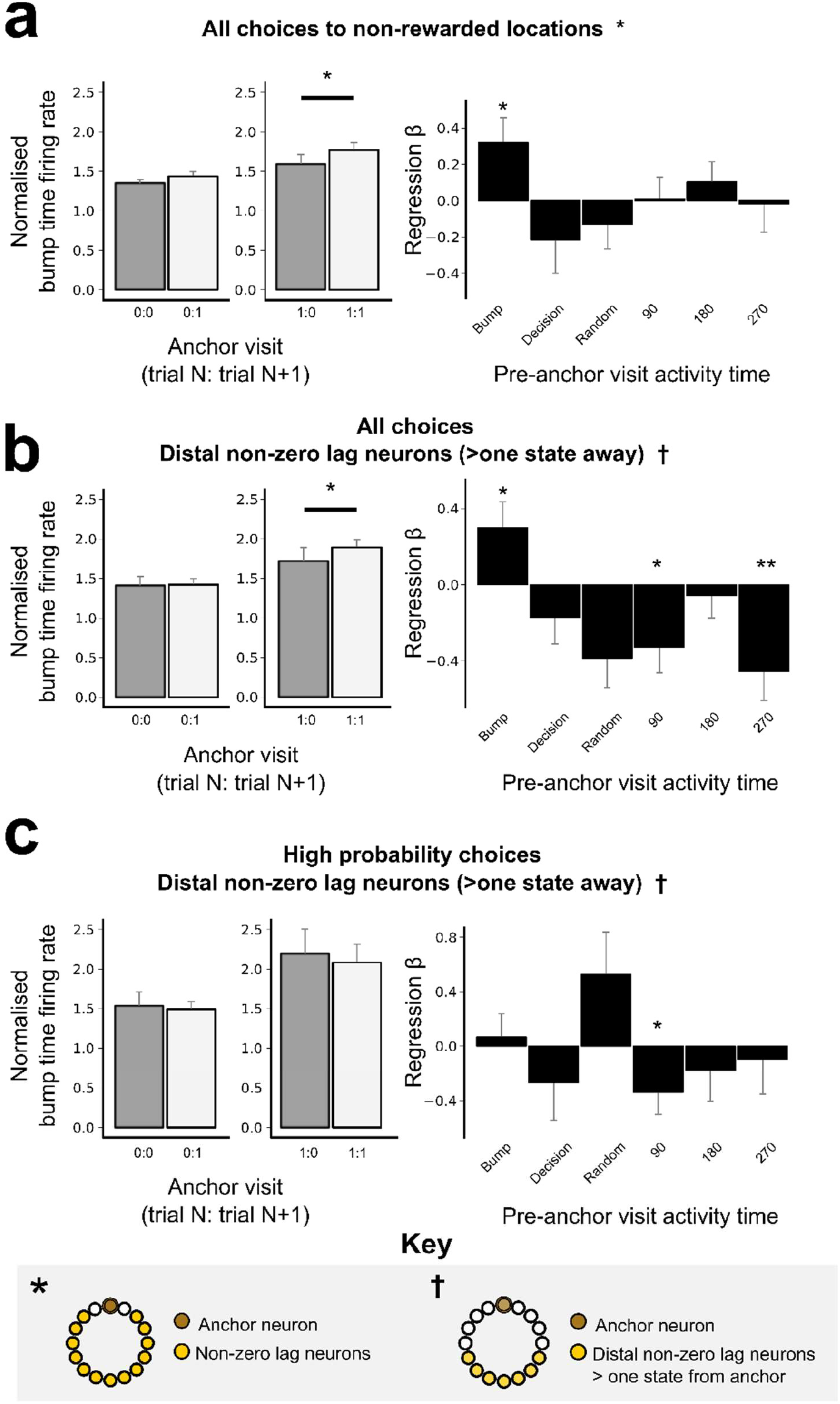
a) Left: Prediction of behaviour to intermediate (non-rewarded) locations i.e. excluding choices to reward locations. Normalised firing rates of neurons during their “bump time”: i.e. the lag at which they are active relative to the anchor. Bump time activity is higher before visits to the neuron’s anchor in trial N+1 whether the anchor was not visited in trial N (left) or when it was visited in trial N (right). Wilcoxon tests: Anchor not visited in trial N: n=76 sessions, statistic=1093, P=0.055. Anchor visited in trial N: n=71 sessions, statistic=880, P=0.023. In addition, an ANOVA on all data (N=71 sessions) showed a main effect of Past: F=11.0 P=0.001, df1=1, df2=70, a trend towards an effect of Future: F=2.74 P=0.103, df1=1, df2=70 and no Past x Future interaction F=0.001 P=0.973, df1=1, df2=70. Right: Regression coefficients were significantly positive for the bump time but not all other control times. T tests against 0: “bump time”: N=76 sessions, statistic=2.32, P=0.023, df=75; “decision time”: N=76, statistic=−1.17, P=0.246, df=75; “random time” N=76, statistic=−0.702, P=0.485, df=75; “90 degree shifted time” N=76, statistic=0.082, P=0.935, df=75; “180 degree shifted time” N=76, statistic=0.956, P=0.342, df=75; “270 degree shifted time” N=76, statistic=−0.112, P=0.911, df=75. b) Prediction of behaviour using neurons anchored at distal points (one state away or more) from the anchor. Normalised firing rates of neurons during their “bump time”: i.e. the lag at which they are active relative to the anchor. Bump time activity is not higher before visits to the neuron’s anchor in trial N+1 when the anchor was not visited in trial N (left) but was higher before visits to anchor in trial N+1 when the anchor was visited in trial N (right). Wilcoxon tests: Anchor not visited in trial N: n=74 sessions, statistic=1130, P=0.225. Anchor visited in trial N: n=66 sessions, statistic=709, P=0.011. In addition, an ANOVA on all data (N=66 sessions) showed a main effect of Past F=16.9 P=1.1×10^−4^, df1=1, df2=65, no main effect of Future F=1.49 P=0.226, df1=1, df2=65 and no Past x Future interaction F=0.006 P=0.939, df1=1, df2=65. Regression coefficients were significantly positive for the bump time but not all other control times. T tests against 0: “bump time”: N=74 sessions, statistic=2.20, P=0.031, df=73; “decision time”: statistic=−1.24, P=0.216, df=73; “random time” N=74, statistic=−1.86, P=0.067, df=73; “90 degree shifted time” N=74, statistic=−2.43, P=0.017, df=73; “180 degree shifted time” N=74, statistic=−0.479, P=0.633, df=73; “270 degree shifted time” N=74, statistic=−2.98, P=0.004, df=73. c) Prediction of behaviour using neurons anchored at distal points (one state away or more) from the anchor for only the high probability choices (choices that animals made with a probability in top half of maze transition probabilities during pre-task exploration). Normalised firing rates of neurons during their “bump time”: i.e. the lag at which they are active relative to the anchor. Bump time activity was indistinguishable before visits to the neuron’s anchor compared to non visits in trial N+1 whether the anchor was not visited in trial N (left) or when it was visited in trial N (right). Wilcoxon tests: Anchor not visited in trial N: n=62 sessions, statistic=808, P=0.323. Anchor visited in trial N: n=50 sessions, statistic=622, P=0.886. In addition, an ANOVA on all data (N=50 sessions) showed a main effect of Past F=10.3 P=0.002, df1=1, df2=49, no main effect of Future F=0.249 P=0.620, df1=1, df2=49 and no Past x Future interaction F=2.09 P=0.155, df1=1, df2=49. Regression coefficients were not significant for the bump time. T tests against 0: “bump time”: N=62 sessions, statistic=0.40, P=0.694, df=61; “decision time”: N=62, statistic=−0.98, P=0.331, df=61; “random time” N=62, statistic=1.70, P=0.095, df=61; “90 degree shifted time” statistic=−2.11, P=0.039, df=61; “180 degree shifted time” N=62, statistic=−0.785, P=0.435, df=61; “270 degree shifted time” statistic=−0.379, P=0.706, df=61. All error bars represent the standard error of the mean

## Methods

### Animals

Experiments used adult male C57BL/6J mice (n = 11; Charles River Laboratories). Animals were run in 2 cohorts of 4 and 1 cohort of 3, and preselected based on criteria outlined in the Behaviour section below. Animals were housed with their littermates up until the start of the experiment with free access to water in a dedicated housing facility with a 12-h light/12-h dark cycle (lights on at 7:00). Food was available *ad libitum* throughout the experiments, and water was available *ad libitum* before the experiments (see below). Mice were 2–5 months old at the time of testing. Experiments were carried out in accordance with Oxford University animal use guidelines and performed under UK Home Office Project Licence P6F11BC25.

### Behavioural training

#### Definitions

##### The ABCD paradigm

A set of tasks where subjects must find a sequence of four **reward locations** (termed ***a,b,c*** and ***d***) in a 3×3 grid maze that repeat in a loop. Once the animal receives reward ***a*** the next available reward is in location ***b*** …etc once the animal gets reward in location ***d*** then reward ***a*** becomes available again, hence creating a repeating loop.

##### Location

Where the animal is in the physical maze

The animal could be at a **node**: i.e. one of the circular platforms where reward could be delivered: coded 1-9 as shown below - they could also be at an **edge** which is a bridge between nodes. (Figure 1a)

##### Task

An example of the ABCD loop with a particular arrangement of reward locations (e.g. 1-9-5-7; reward ***a*** is in location 1; reward ***b*** in location 9 …etc; Figure 1a)

##### Session

An uninterrupted block of trials of the same task. We used 20 minute sessions. Note that subjects could be exposed to 2 or more sessions of the same task on a given day. Animals were allowed to complete as many trials as they could in those 20 minutes. Animals were removed from the maze at the end of a session and either placed back in their home-cage or into a separate enclosed rest/sleep box.

##### Trial

A <u>single</u> run through an entire ABCD loop for a particular task, starting with reward in location ***a*** and ending in the next time the animal gets reward in location ***a*** again (e.g. trial 12 of a task with the following configuration: 1-9-5-7 starts with the 12th time the animal gets reward in location 1 and ends with the 13th time animal gets reward in location 1)

##### State

The time period between an animal receiving reward in a particular location and receiving reward in the next rewarded location. State **A** (upper case) starts when animal receives reward ***a*** (lower case italicised) and ends when animal receives reward ***b***…state **D** starts when animal gets reward ***d*** and ends when animal gets reward ***a*** …etc

##### Transition

A generalised definition of state. e.g. progressing from ***a*** to ***b*** is a transition, and from ***c*** to ***d*** is also a transition.

##### Goal-progress

How much progress an animal has made between rewarded locations as a percentage of the time taken between them. Unless otherwise stated, we operationally divide this into early, intermediate and late goal-progress - which correspond to ⅓ increments of the time taken to get from one reward location to the next: e.g. if the animal takes 9 seconds between one reward and another, then early goal-progress spans the first 3 seconds, intermediate goal-progress the next 3 seconds and late goal-progress the last 3 seconds. This would scale with the length of time it takes for the animal to complete a single state: e.g. if it takes 15 seconds between two rewards, each of the goal-progress bins would be 5 seconds long. In the ABCD paradigm goal-progress repeats 4 times because there are 4 rewards (so there will be an early goal-progress for reward ***a***, and early goal-progress for reward ***b*** …etc).

##### Choice

We use this to refer to one-step choices in the maze. At every node in the maze the animal has a choice of 2 or more immediately adjacent nodes to visit next. For example, when in node 1 the animal could choose to move to node 2 or node 4 (Figure 1a).

#### Pre-selection

A total of 11 mice, across 3 cohorts, were used for experiments. For each cohort, 3-4 Mice were pre-selected for the experiment from 10-20 mice based on the following criteria:

1. Weight above 22g
2. No visible signs of stress upon first exposure to the maze in the absence of rewards. Stress was evidenced by thigmotaxis or defecation in a 20 minute exploration session with no rewards delivered
3. On a partially connected version of the maze with only 6 accessible ports out of 9, Animals learned that poking in wells delivered water reward and that after gaining reward they must go to another port. Animals that obtained 40 or more rewards per 20 minute session were taken forward to pretraining.
4. Final selection

For a given cohort, if more than 4 animals satisfied these criteria, animals that explored the maze with the highest entropy were selected (see “Behavioural Scoring” below for entropy calculation).

#### Habituation/Pretraining

After at least one week of post-surgery recovery (see “Surgeries” section below), animals were placed on water restriction. Animals were maintained at a weight of 88-92% of their baseline weight, which was calculated before water restriction but after implantation and recovery from the surgery (see below under “Surgeries”). This is to ensure that they remained motivated to collect water rewards during the task but not overly so, as excessive motivation is known to negatively affect model-based performance^44^. Animals were habituated to being tethered to the electrophysiological recording wire while moving on the maze, as well as in sleep boxes for at least 3 days prior to the start of the experiment. During this period, animals were reintroduced to a partially connected maze (only 6 out of the 9 ports available, and not all connected) while tethered to the electrophysiology wire, where water reward was delivered if the animal poked its nose at any port. Reward drops were only available once the animal poked its nose at the port. At this stage, there was no explicit task structure, except that once reward was obtained at one port, animals had to visit a different port to gain further reward (i.e. exactly as in step 3 of pre-selection criteria above but while implanted and tethered). Thus, animals (re)learned that poking in wells delivered reward and that after gaining reward they must go to another port. Animals were transitioned to the task when they obtained >40 rewards in a 20 minute session. Note that 3 animals (from cohort 1) were not implanted and so for these the rewarded sessions for pre-selection and pre-training were one and the same (see section below under “Numbers” for more details on mouse cohorts). The volume of water delivered during pretraining/preselection was higher than training, typically 10-15 µL.

#### Training

Animals navigated an automated 3×3 grid maze in search of rewards (Figure 1a), controlled using pyControl^51^. Water rewards (3-4 µL) were presented sequentially at 4 different locations. Animals had to poke in a given reward port, breaking an infrared beam that triggered the release of the reward drop in this well. After reward ***a*** was delivered, reward was obtainable from location ***b***, but only after the animal poked in the new location. Once the animal received reward in locations ***a***, then ***b***, then ***c*** and then ***d***, reward ***a*** becomes available again, thus creating a repeating loop. Animals have 20 minutes to collect as many rewards as possible and no time outs are given if they make any mistakes. They are then allowed at least a 20 minute break away from the maze (in the absence of any water) before starting a new session. For each session, animals were randomly entered from a different side of the square maze, using custom made electromagnetic field shielding curtains (https://www.electrosmogshielding.co.uk/product.asp?P_ID=650&CAT_ID=104). This was to ensure that all sides of the maze are equivalent in terms of being entry and exit points from the maze, thereby minimising any place preference/aversion and minimising the use of different sides as orienting cues. One cue card was placed high up (at least 50 cm vertically from the maze) on one corner of the maze to serve as an orienting cue. No cues were visible at head level.

While all locations were rewarded identically, a brief pure tone (2 seconds at 5 kHz) was delivered when the animal consumed reward ***a***. This ensured that task states were comparable across different task sequences. White noise was present throughout the session to avoid distraction from outside noises.

Task configurations (i.e. the sequence of reward locations) were selected pseudo-randomly for each mouse, while satisfying the following criteria:

1. The distance between rewarded locations in physical space (number of steps between rewarded locations) and task space (number of task states between rewarded locations) were orthogonal for each mouse (Extended data Figure 1a)
2. The task can’t be solved (75% performance or more) by moving in a clockwise or anti-clockwise circle around the maze.
3. The first two tasks have location 5 (the middle location) rewarded - this is to ensure the first tasks the animals are exposed to can’t be completed by circling around the outside N times to collect all rewards. (Note that all “late” tasks and those used in electrophysiological recordings are not affected by this criterion).
4. Consecutive tasks don’t share a transition (i.e. two or more consecutively rewarded locations)
5. Chance levels are the same for all task transitions (***ab***, ***bc***, ***cd***, ***da***), and control transitions ***ca*** and ***ba*** transitions - whether determined analytically by assuming animals diffuse around the maze or empirically by using animal-specific maze-transition statistics from an exploration session before any rewards were delivered on the maze (see “Behavioural Scoring” below for chance-level calculations).

For the first 10 tasks, animals were moved to a new task when their performance reached 70% (i.e. took one of the shortest spatial paths between rewards for at least 70% of all transitions) on 10 or more consecutive trials or if they plateaued in performance for 200 or more trials. For these first 10 tasks, animals were given at most 4 sessions per day, either all of the same task or, when animals reached criteria, two sessions of the old task and two sessions of the new task. From task 11 onwards, animals learned 3 new tasks a day with the first task being repeated again at the end of each day giving a total of 4 sessions with the pattern X-Y-Z-X’.

### Surgeries

Subjects were taken off water restriction 48 hours before surgery and then anaesthetised with isoflurane (3% induction, 0.5–1% maintenance), treated with buprenorphine (0.1 mg kg−1) and meloxicam (5 mg kg−1) and placed in a stereotactic frame. A silicon probe mounted on a microdrive (Ronal Tool) and encased in a custom made recoverable enclosure (ProtoLabs) was implanted into mFC (AP: 2.00, ML: −0.4, DV: −1.0), and a ground screw was implanted above the cerebellum. AP and ML coordinates are relative to bregma while DV coordinates are relative to the brain surface. Mice were given additional doses of meloxicam each day for 3 days after surgery and were monitored carefully for 7 days after surgery and then placed back on water restriction 24 hours before pretraining. At the end of the experiment, animals were perfused; and the brains were fixed-sliced and imaged to identify probe locations (Extended data figure 2a).

### Electrophysiology, spike sorting and behavioural tracking

Cambridge NeuroTech F-series 64 silicon channel probes (6 shanks spanning 1 mm arranged front-to-back along the anterior posterior axis) were used for all recordings. To record from the mFC, we lowered the probe ~100 µm during the pre-habituation period to reach a final DV position of between −1.3 and −1.5 mm below the brain surface (i.e. between −2.05 and −2.25 mm from bregma). This places most channels in the prelimbic cortex (http://labs.gaidi.ca/mouse-brain-atlas/). Neural activity was acquired at 30 kHz with a 32-channel Intan RHD 2132 amplifier board (Intan Technologies) connected to an OpenEphys acquisition board. Behavioural, video and electrophysiological data were synchronised using sync pulses output from the pyControl system. Recordings were spike sorted using Kilosort^52^, versions 2.5 and 3, and manually curated using phy (https://github.com/kwikteam/phy). Clusters were classified as single units and retained for further analysis if they had a characteristic waveform shape, showed a clear refractory period in their autocorrelation and were stable over time. We performed tracking of the mice in the video data using DeepLabCut^53^, a Python package for marker-less pose estimation based in the TensorFlow machine learning library. Positions of the back of a mouse’s head in x, y pixel coordinates were converted to region of interest information (which maze node or edge the animal is in for each frame) using a set of binary masks defined in ImageJ that partition the frame into its sub components.

### Behavioural Scoring

Performance was assessed by quantifying the percentage of transitions where animals took one of the shortest available routes between two rewards. We also quantified the path length taken between rewards and divided this by the shortest length to give the “relative path distance” covered per transition.

When using “percentage of shortest path transitions” as a criteria, chance levels were calculated either analytically or empirically. Analytical chance levels were calculated by assuming a randomly diffusing animal and calculating the chances the animal will move from node X to node Y in N steps. Empirical chance levels were calculated by using the location-to-location transition matrix recorded for each animal in the Exploration session before any exposure to reward on the maze. When using the “relative path distance” measure, chance was calculated empirically using the mean relative path distance for transitions in the first trial averaged across the first 5 tasks.

Correct transition entropy (The animal’s entropy when taking the shortest route between rewards) was calculated for transitions where there was more than one shortest route between rewards. We calculated the probability distribution across all possible shortest paths for a given transition and calculated entropy as follows:

**Entropy = ∑pk.log_x_(pk)**

Where **x** is the logarithmic base which is set to the number of shortest routes and **pk** is the probability of each transition. Thus an entropy of 1 signifies complete absence of a bias for taking any one path and an entropy of 0 means only one of the paths is taken (i.e. maximum stereotypy).

### Activity Normalisation

We aimed to define a task space upon which to project the activity of the neurons. To achieve this, we aligned and normalised vectors representing neuronal activity and maze location to the task states. Activity was aligned such that the consumption of reward ***a*** formed the beginning of each row (trial) and consumption of the next reward ***a*** started a new row. Normalisation was achieved such that all states were represented by the same number of bins (90) regardless of the time taken in each state. Thus, the first 90 bins in each row represented the time between rewards ***a*** and ***b***, the second between ***b*** and ***c,*** the third between ***c*** and ***d*** and the last between ***d*** and ***a***. We then computed the averaged neuronal activity for each bin. Thus the activity of each neuron was represented by an Nx360 matrix, where N is the number of trials and 360 bins represent task space for each trial. This activity was then averaged by taking the mean across trials, and smoothed by fitting a Gaussian kernel (sigma=10 degrees). To avoid edge effects when smoothing, the mean array was concatenated to itself three times, then smoothed, then the middle third extracted to represent the smoothed array. To reflect the circular structure of the task, the mean and standard error of the mean of this normalised and smoothed activity were projected on polar plots (e.g. Figure 2b-d).

### Tuning to basic task variables

#### Generalised linear model

To assess the degree to which mFC neurons are tuned to task space, we used a linear regression to model each neuron’s activity. Specifically, we aimed to quantify the degree to which goal-progress and location tuning of the neurons is consistent across tasks and states. For this we used a leave-one-out cross-validation design: we divided all tasks into the time periods spanned by each of the 4 states and used all data except one Task/State combination to train the model. The remaining task/state (e.g. task 3, state B) was used to test the model. This was repeated so that each task/state combination had been left out as a test period once. The training periods were used to calculate mean firing rates for 5 levels of goal-progress relative to reward (5 goal-progress bins) and each maze location (9 possible node locations). Edges were excluded from analyses since they are systematically not visited at the earliest goal-progress bin. The mean firing rates for goal-progress and place from the training sessions were used as (separate) regressors to test against the binned firing rate of the cell in the test data (held out task-state combination). To assess the validity of any putative task tuning, a number of potentially confounding variables were added to the model. These were: acceleration, speed, time from reward, and distance from reward. This procedure was repeated for all Task/State combinations and a separate regression coefficient value was calculated for each. To assess significance per neuron, we repeated the regression but with random circular shifts of each neuron’s activity array and computed regression coefficient values for each iteration (100 iterations) and then used the 95th percentile of this distribution as the regression coefficient threshold. Neurons with an actual regression coefficient value higher than this threshold were considered to be tuned for this variable. Two proportions Z-tests were performed to assess whether the proportion of neurons with significant regression coefficient values for a given variable were statistically higher than a chance level of 5%.

#### State tuning

For state tuning, we first wanted to test whether neurons were tuned to a given state in a given task. We therefore analysed state tuning separately from the GLM above, which explicitly tests for the consistency of tuning across tasks. Instead, we used a z-scoring approach. First we took the peak firing rate in each state and trial, giving 4 values per trial: i.e. a *maximum activity* matrix with dimensions Nx4 where N is the number of trials. Then we z-scored each row of this maximum activity matrix (i.e. giving a mean of 0 and standard deviation of 1 for each trial). We then extracted the z-scores for the preferred state across all N trials and subsequently conducted a T test of this array against 0. Neurons with a P value of < 0.05 for a given task were taken to be state tuned in that task.

##### Neuronal generalisation

To assess whether individual neurons maintained their state preference across tasks we quantified the angle made between a neuron in one task and the same neuron in another task. Only state-tuned neurons were used in these analyses. To ensure we captured robustly state tuned neurons, we restricted analyses to neurons state tuned in more than 1/3rd of the recorded tasks. This subsetting is used throughout the manuscript where state tuned cells are investigated. Quantifying the angle between neurons was achieved by rotating the neuron in task Y by 10 degree intervals and then computing the Pearson correlation between this rotated firing rate vector and the mean firing rate vector in task X. Using this approach, we found for each neuron the rotation that gave the highest correlation. For cross-task comparisons we calculated a histogram of the angles across the entire population and averaged this across both comparisons (X vs Y and X vs Z). Within task histograms were computed by comparing task X to task X’ (Figure 3b). To compute the proportion of neurons that generalised their state tuning, we found the maximum rotation across both comparisons (X vs Y and X vs Z). We then set 45 degrees either side of 0 rotation across all tasks as the generalisation threshold. Because this represents ¼ of the possible rotation angles, chance level is equal to ¼^m^ where m is the number of comparisons. When calculating generalisation across one comparison, chance level is therefore 25%, whereas when two comparisons are taken, the chance level is 1/16 (6.25%).

Generalisation could also be expressed at the level of tuning relationships between neurons. For example two neurons that are tuned to A and C in one task could then be tuned to B and D in another, thereby maintaining their task-space angle (180 degrees) but remapping in task space across tasks. To test for this, we computed the tuning angle *between* pairs of neurons and assessed how consistent this was across tasks. This angle was computed by rotating one neuron by 10 degree intervals and calculating the Pearson correlation between the mean firing vector of neuron k and the rotated firing vector for neuron j. The rotation with the highest Pearson correlation gave the between-neuron angle (Figure 3c). Thus, we compared the angle between a pair of neurons in task X to the same between-neuron angle in tasks Y and Z. Again histograms were averaged across both comparisons (X vs Y and X vs Z) for cross-task histograms while within-task histograms were computed by comparing task X to task X’ (Figure 3c). To compute the proportion of neuron pairs that were coherent across tasks, we found the maximum rotation of the angle between each pair across both comparisons (X vs Y and X vs Z). We then set 45 degrees either side of 0 rotation across all tasks as the coherence threshold. Because this represents ¼ of the possible rotation angles, chance level is equal to ¼^m^ where m is the number of comparisons and therefore=25% for one comparison and 1/16 (6.25%) for two comparisons.

To assess whether the mFC population was organised into modules of coherently rotating neurons, we used a clustering approach. The first step was to take the *maximum* difference in pairwise, between-neuron angles (across all comparisons) and convert this into a maximum circular distance (1-cosine(angle)) thereby generating a distance matrix reflecting coherence relationships between neurons (incoherence matrix). The second step is to compute a low dimensional embedding of this incoherence matrix, using t-distributed stochastic neighbour embedding (t-SNE; using the TSNE function of scikit learn manifold library, with perplexity=5). Finally, the third step is to use hierarchical clustering on this embedded data (using the AgglomerativeClustering function of scikit learn cluster library, with distance threshold=300). This procedure sorts neurons into clusters reflecting coherence relationships between neurons. We quantified the degree of clustering by computing the Silhouette Score for the clusters computed in each recording day:

### Silhouette Score=(b-a)/max(a,b)

Where a=mean intracluster distance, b=mean intercluster distance. We repeated the same procedure but for permuted data, where state tuning in task X and goal-progress tuning in all tasks was identical to the real data but the state preference of each neuron remapped randomly across tasks. This allowed us to compare the Silhouette Scores for the real and permuted data (Figure 3d). To visualise clusters and the tuning of neurons within them in the same plot, we plotted some example neurons from a single recording day where the x and the y axes represented state tuning and the y axis arranged neurons based on their cluster id (the ordering along the z axis is arbitrary; Figure 3e).

#### Mnemonic task space tuning

The Task structured memory buffers (SMBs) model predicts the existence of neurons that maintain an invariant task space lag from a particular anchor representing a behavioural step, regardless of the task sequence. Concretely, behavioural steps are conjunctions of goal-progress (operationally divided into early, intermediate or late) and place (nodes 1-9). To test this prediction, we used three complementary analysis methods. All of these analyses were conducted on data where two recording days were combined and spike-sorted concomitantly, giving a total of 6 unique tasks per animal (with two exceptions that had 4 and 5 tasks each; see exclusions under the “Numbers” section below). For all of these analyses, only state tuned neurons were used (see “Tuning to basic task variables”).

##### Method 1: Model fitting

For each neuron we computed a regression model that described state-tuning activity as a function of all possible combinations of goal-progress/place and all task lags from each possible goal-progress/place. Thus a neuron could fire at a particular goal-progress/place but also at a particular lag in task space from this goal-progress/place. We used an Elastic Net (using the ElasticNet function from the scikit learn linear_model package) that included a regularisation term which was a 1:1 combination of L1 and L2 norms. The alpha for regularisation was set to 0.01. A total of 9×3×12 (312) regressors were used for each neuron, corresponding to 9 locations, 3 goal-progress bins (so 27 possible goal-progress/place anchor points) and 12 lags in task space from the anchor (4 states x 3 goal-progress bins). We trained the model on 5 (training) tasks and then used the resultant regression coefficients to predict the activity of the neuron in a left out (test) task. Both training and cross-validation were only done in the preferred goal-progress of each neuron to ensure our prediction results are due to state preference and not the strong effect of goal-progress preference (Figure 2 e,f). For non-zero-lag neurons, we only used state-tuned neurons with all of the three highest regression coefficient values at non-zero lag from an anchor (lag from anchor of 30 degrees or more for Figure 5b bottom; 90 degrees (one state) or more for Extended data figure 5b) in the training tasks. Also, for non-zero lag neurons, we only use regression coefficient values either 30 degrees (Figure 5b) or 90 degrees (Extended data figure 5b) either side of the anchor point to predict the state tuning of the cells. This ensured that the prediction was only due to lagged activity and not direct tuning of the neurons to the goal-progress/place conjunction.

##### Method 2: Lagged spatial similarity

To detect putative mnemonic task space neurons, we calculated spatial tuning to where the animal was at different task lags in the past (Figure 5c). While spatial neurons should consistently fire at the same locations(s) at zero lag, neurons that track a memory of the goal-progress/place anchor will instead show a peak in their cross-task spatial correlation at a non-zero task lag in the past (Figure 5c). To quantify this effect, we used a cross-validation approach, using all tasks but one to calculate the lag at which cross-task spatial correlation was maximal, and then measuring the Pearson correlation between the spatial maps in the left out task and the training tasks at this lag (Figure 5d). To account for the strong goal-progress tuning of the neurons, all maps were computed in each neuron’s preferred goal-progress.

##### Method 3: Single anchor alignment

This approach assumes each neuron can only have a single goal-progress/place anchor and quantifies the degree to which task-space lag for this neuron is conserved across tasks. We fitted the anchor by choosing the goal-progress/place conjunction which maximises the correlation between lag-tuning-curves in all but one (training) tasks, and again used cross-validation by assessing whether this anchor leads to the same lag tuning in the left out (test) task. The fitting was conducted by first identifying the times an animal visited a given goal-progress/place and sampling 360 bins (1 trial) of data starting at this visit, then averaging activity aligned to all visits in a given task, and smoothing activity as described above (Under “Activity Normalisation”). This realigned activity is then compared across tasks to compute the angle (θ) between the neuron’s mean aligned/normalised firing rate vector across tasks. This involves essentially doing all the steps for the “Neuronal generalisation analysis” but for the anchor-aligned activity instead of state-A-aligned activity. This was done for all possible task combinations and then a distance matrix (M) was computed by taking distance=1-cosine(θ). This distance matrix M has dimensions Ntraining_tasks x Ntraining_tasks x Nanchors (typically 5×5×27 as there are usually 6 tasks, meaning 5 training tasks are used, and 3×9 possible anchors corresponding to 3 possible phases (early, intermediate and late) and 9 possible maze locations). The distance can then be averaged across all comparisons to find the mean distance between all comparisons for a given anchor for a given training task - generating a mean-distance matrix (M_mean). This M_mean matrix has the dimensions Ncomparisons x Nanchors; where Ncomparisons= Ntraining_tasks – 1; typically this will be 4 × 27). The entry with the minimum value in this N_mean matrix gives the combination of training task and goal-progress/place anchor that best aligns the neuron – the training task selected is referred to as the reference task. Next, the neuron’s mean activity in the test task is aligned to visits to the best anchor calculated from the training tasks. This allows calculating how much this aligned activity array has remapped relative to the aligned activity in the reference training task - if it has remapped by 0 degrees or close to 0 degrees (within a 45 degree span either side of zero) then the neuron is spatially anchored (i.e. maintains a consistent angle with its anchor across tasks). For a given test-train split, we computed a histogram of the angles across all the neurons and then we averaged the histograms across all test-train splits to visualise the overall distribution of angles between training and test tasks (e.g. Figure 5f). To quantify the degree of alignment further, we measured the correlation between the anchor-aligned activity of neurons in the test task versus reference training task; only considering activity of neurons in their preferred goal-progress bin (Figure 5g). The neuron’s lag from its anchor was identified by finding the lag at which anchor-aligned activity was maximal. This lag is used below for “Predicting Behavioural Choices”.

For per mouse effects reported in Extended data figure 5, we tested whether the number of mice with a mean cross-validated correlation above 0 is higher than chance, chance level being a uniform distribution (50:50 distribution of per mouse correlation means above and below zero). We used a one-tailed binomial test against this chance level. All 5 mice need to have mean positive values for this test to yield significance.

#### Predicting Behavioural Choices

The SMB model proposes that behavioural choices should be predictable from bumps of activity along specific memory buffers long before an animal makes a particular choice. By “choice” here we mean a decision to move from one node to one of the immediately adjacent nodes on the maze (e.g. from location 1 should I go to location 2 or 4?; Figure 6a). To test whether these choices are predictable from distal neuronal activity we used a Logistic regression model. For this analysis we used only consistently anchored neurons, that is neurons that had the same anchor and same lag to anchor in at least half of the tasks. This relied on the single-anchor analysis (Figure 5e-g). Furthermore, to avoid contamination of our results due to simple spatial tuning, we only used neurons with activity lagged far from their anchor (at least 30 degrees in task space either side of the anchor e.g. Figure 6b,d,e,f; and this was also repeated for lags of at least 90 degrees (one whole state) either side of the anchor: e.g. Figure 6g and Extended data 6b,c). We measured the activity of a given neuron during its “bump” time, that is the time at which a neuron is lagged relative to its anchor. Precisely, this is the mean firing rate from a period starting with the lag time from the anchor and ending 30 degrees forward in task space from that point (i.e. 1/3rd of a state). This mean activity was inputted on a trial by trial basis every time the animal was at a goal-progress/place conjunction that was one step before the goal-progress/place anchor in question (e.g. if the anchor is at early goal-progress in place 2, the possible goal-progress/places before this are: late goal progress in place 1, late goal-progress in place 3 and late goal-progress in place 5; see maze structure in Figure 1a). We used this activity to predict a binary vector that has 1s when the animal visits the anchor and 0 when the animal could have visited the anchor (i.e. was one step away from it) but did not chose to visit the anchor. To remove confounds due to the autocorrelated previous behavioural choices, we added previous choices up to 5 trials in the past into the regression model. Furthermore, to assess whether any observed prediction was specific to the “bump” time as predicted by the SMB model, we repeated the logistic regression for other times (random times, decision time (30 degrees from the potential anchor visit) and times shifted by 90, 180 and 270 degrees from the bump time). We further repeated this regression only for visits to non-zero goal-progress anchors (i.e. non-rewarded locations) and also only taking neurons that are more distal from their anchor (i.e. at least 90 degree separation either side of the anchor). We also conducted this analysis separately for high probability choices (choices that were in the top 50% of the maze transition probability distribution calculated in a baseline exploration session before any reward or task was ever experienced on the maze; e.g. Figure 6e) and low probability choices (bottom 50% of the same transition probability distribution; e.g. Figure 6f). This allowed us to test whether frontal neurons generally predict all choices or more specifically predict choices which animals were not already predisposed to make.

#### Sleep/Rest analysis

To investigate the internal organisation of task-related mFC activity we recorded neuronal activity in a separate enclosure containing bedding from the animal’s home cage but no reward or task-relevant cues. Animals were pre-habituated to sleep/rest in these “sleep boxes” before the first task began. We measured neuronal activity across sleep/rest sessions both before any tasks on a given day and after each session. The first (pre-task) sleep/rest session was 1 hour long, inter-session sleep/rest sessions were 20 minutes long and the sleep/rest session after the last task was 45 minutes long. All sessions except the first sleep session were designated as “post-task” sleep sessions.

We wanted to assess whether the memory buffers were organised by the task structure, which in our case is a ring. In other words, do neurons internally maintain this ring-like organisation even in the absence of externally structured sensory/behavioural input? Activity was binned in 250 ms bins and cross-correlations between each pair of neurons were calculated using this binned activity. We then regressed the awake angle difference between pairs of neurons sharing the same anchor against this sleep cross-correlation. This angle was taken from the first task on a given day for pre-task sleep, and from the task immediately before the sleep session for all post-task sleep sessions. The idea is that neurons closer to each other in a given neuronal state-space should be more likely to be coactive within a small time window compared to neurons farther apart. To control for place and goal-progress tuning, we added the spatial map correlation and circular goal-progress distance as co-regressors. Thus we assessed the degree to which the regression coefficients were negative (i.e. smaller distances correlate with higher coactivity).

We first sought to identify whether the neurons were functionally organised into rings or delay lines. For this we measured the forward distance between pairs of co-anchored neurons in reference to their anchor. If neurons are internally organised on a line, then the larger the forward distance between a pair of neurons the further away two neurons are from each other in neuronal state space, and hence the less coactive they will be (Figure 7a). If neurons are arranged on a ring, forward distance should also positively correlate with neuronal state space distance for pairs of neurons separated by less than 180 degrees in task space. Beyond 180 degrees, forward distance should instead negatively correlate with neuronal state space distance, as neurons circle back towards each other (Figure 7a). In this situation, circular distance, not forward distance, would be the best description of the neuronal state space. Thus, for a delay line, the higher the forward distance beyond 180 the further the neurons should be from one another, and hence the coefficient for the regression between forward distance and coactivity should be negative (Figure 7a). For a ring, the coefficient value for forward distance against coactivity should conversely be positive. This pattern inverts when considering the circular distance: positive coefficient for delay line and negative coefficient for rings. We used a linear regression to compute the regression coefficients for circular and forward distances in the same regression for *all* neuron pairs (Figure 7b). To control for place and goal-progress tuning, we again added the spatial map correlation and circular goal-progress distance as co-regressors. We also used a linear regression to compute this coefficient for only the circular distance for pairs of neurons separated by 180 degrees or more, while again co-regressing spatial correlations and goal-progress distances (Figure 7c). We further analysed whether neurons that shared the same anchor showed stronger state-space relationships than those that have different anchors. For this we restricted our analysis to consistently anchored neurons, that is neurons that had the same anchor and same lag to anchor in at least half of the tasks. We conducted the regression of circular distance against sleep cross-correlation either for pairs of neurons that share the same anchor, or those that have different anchors (Figure 7d). As before, we co-regressed spatial correlations and goal-progress distances.

Note that, arguably, the last analysis (Figure 7d) could be seen as sufficient to indicate a ring-like structure. However, this is only true when assuming a uniform distribution of pairwise angles, which we know not to be the case given the overrepresentation of neurons with a small lag from the anchor (Extended data Figure 5c), thus necessitating the first two analyses (Figure 7b,c).

#### Numbers

##### Animals

11 animals in total were used for behavioural recordings across 3 separate cohorts conducted by 2 different experimenters (AH and ME) - 4 of these animals only completed 10 tasks as part of the first cohort and the remaining 7 completed 40 tasks Of the 11 animals, 5 animals in total were used for electrophysiological recordings, the remaining 6 animals are accounted for below:

- 3 animals (in cohort 1) were not implanted at any point
- 1 animal was implanted with silicon probes but was part of the first cohort so did not get to the 3 task days (i.e. only completed the first 10 tasks)
- 2 animals were implanted but their signal was lost before the 3 task days

Exclusions: No animals were excluded from analyses: All animals (11) were included in the behavioural analyses, and all animals for which there was an electrophysiological signal by the 3 task days (5) were included in the electrophysiological analyses.

##### Electrophysiological recording days

All electrophysiological data was obtained from days where animals completed 3 tasks per day (cohorts 2 and 3) and in late tasks (tasks 21-40) where animals showed robust evidence for having learned the abstract task structure (Figure 1). 33 recording days were spike sorted as individual days and were used in Figure 2 and Figure 3. In 23 of those 32 days, animals completed more than 2 trials in the last session (the repeat of task X), hence fewer neurons appear in analyses comparing session X with session X’ (Figure 3b,c and Extended data figure 3b,c). 14 recording days were merged to create double-days, allowing tracking of neuronal activity across up to 6 tasks. Although 6 tasks were always attempted across each double day, of these 14 double days, 12 spanned 6 tasks, 1 spanned 5 tasks and 1 spanned 4 tasks. This was due to file corruption (1 task) or animals lacking motivation (less than 2 trials completed per task; 2 tasks). Double days were used in Figures 3,5,6 and 7. Only single days were used in Figure 2.

Data was obtained from all possible double-days from late tasks (tasks 21-40). This means 3 double days per mouse (spanning the last 18 tasks) - except for two mice where each had one double-day where the signal had dropped, and one mouse where 4 double days were used (including a day spanning tasks 20-22) - thus a total of 14 double-days were used.

##### Total number of neurons

We report “neuron-days” - i.e. by summing up each day’s neuron yield below:

1807 total “neuron-days” (i.e. while splitting each double-day into two and summing across days)

1438 separately sorted neuron days (i.e. using a single yield for each double day), of which there were:

- 747 neuron days on single days
- 691 neuron days on double days

Exclusions:

- Recording days with 10 or fewer state-tuned neurons were excluded from analyses of pairwise coherence between neurons.
- Neurons whose mean firing rate during sleep dropped to less than 20% of their awake firing rate were excluded from preplay/replay analysis - this was assumed to be largely a spike sorting artefact.
- Only neurons that were significantly state tuned (neurons with significant state tuning in more than 1/3rd of all tasks) and were sorted across double days were used in Figure 5 - this amounted to 350 neurons - although for the regression analysis, 10 additional neurons were excluded due to having all zero regression coefficient values (caused by the regularisation), giving a total of 340 neurons for this analysis (Figure 5a,b).

## Acknowledgements

We would like to thank Gil Costa for designing the model schematics; Diksha Gupta, Michael Bukwich, Thomas Mrsic-Flogel, Ray Dolan and Masahiro Nakano for valuable input on the manuscript; Raquel Pinacho, Lauren Burgeno, Beatriz Godinho, Marta Blanco Pozo, Veronika Samborska, for guidance on data collection and data preprocessing; All members of the Behrens and Walton labs for their input and support during the project.

ALH is supported by a Wellcome Trust PhD studentship (220047/Z/19/Z), TEJB is supported by a Wellcome Principal Research Fellowship (219525/Z/19/Z) and a Wellcome Collaborator award (214314/Z/18/Z). The Wellcome Centre for Integrative Neuroimaging and Wellcome Centre for Human Neuroimaging are each supported by core funding from the Wellcome Trust (203139/Z/16/Z, 203147/Z/16/Z). JCRW.; is supported by the Sir Henry Wellcome Post-doctoral Fellowship (222817/Z/21/Z). WD is supported by the Gatsby Charitable Foundation. TA is supported by the Wellcome Trust career development award: 225926/Z/22/Z. MEW is supported by a Wellcome Collaborator award (214314/Z/18/Z) and a Wellcome trust SRF (202831/Z/16/Z).

## Author Contributions

ME, JCRW, MEW, TA and TEJB conceptualised the study and designed the ABCD task; ME, ALH and AB performed the surgeries and collected the behavioural and electrophysiological data with input from TA and MEW; ME and ALH analysed and interpreted the data with input from TEJB and TA; ME conceptualised the model with input from TEJB; ME, ALH, JCRW, WD, MW, TA and TEJB elaborated on the core model; ME, ALH, TA and TEJB wrote the manuscript and made the figures with input from all other authors.

## Materials & Correspondence

Correspondence to Mohamady El-Gaby and Tim EJ Behrens.

## Data Availability

All data will be made available upon publication.

## Code Availability

All code will be made available upon publication.

